# Leveraging co-evolutionary insights and AI-based structural modeling to unravel receptor-peptide ligand-binding mechanisms

**DOI:** 10.1101/2024.01.18.575556

**Authors:** Simon Snoeck, Hyun Kyung Lee, Marc W. Schmid, Kyle W. Bender, Matthias J. Neeracher, Alvaro D. Fernández-Fernández, Julia Santiago, Cyril Zipfel

## Abstract

Secreted signaling peptides are central regulators of growth, development, and stress responses, but specific steps in the evolution of these peptides and their receptors are not well understood. In addition, the molecular mechanisms of peptide-receptor binding are only known for a few examples, primarily owing to the limited availability of structural capabilities to few laboratories worldwide. Plants have evolved a multitude of secreted signaling peptides and corresponding transmembrane receptors. Stress-responsive SERINE RICH ENDOGENOUS PEPTIDES (SCOOPs) were recently identified. Bioactive SCOOPs are proteolytically processed by subtilases and are perceived by the leucine-rich repeat receptor kinase MALE DISCOVERER 1-INTERACTING RECEPTOR-LIKE KINASE 2 (MIK2) in the model plant *Arabidopsis thaliana*. How SCOOPs and MIK2 have (co-)evolved, and how SCOOPs bind to MIK2 are however still unknown. Using *in silico* analysis of 350 plant genomes and subsequent functional testing, we revealed the conservation of MIK2 as SCOOP receptor within the plant order Brassicales. We then leveraged AlphaFold-Multimer and comparative genomics to identify two conserved putative SCOOP-MIK2 binding pockets across Brassicales MIK2 homologues predicted to interact with the ‘SxS’ motif of otherwise sequence-divergent SCOOPs. Notably, mutagenesis of both predicted binding pockets compromised SCOOP binding to MIK2, SCOOP-induced complex formation between MIK2 and its co-receptor BRASSINOSTEROID INSENSITIVE 1-ASSOCIATED KINASE 1 (BAK1), and SCOOP-induced reactive oxygen species production; thus, confirming our *in silico* predictions. Collectively, in addition to revealing the elusive SCOOP-MIK2 binding mechanisms, our analytic pipeline combining phylogenomics, AI-based structural predictions, and experimental biochemical and physiological validation provides a blueprint for the elucidation of peptide ligand-receptor perception mechanisms.

**Significance statement:** This study presents a rapid and inexpensive alternative to classical structure-based approaches for resolving ligand-receptor binding mechanisms. It relies on a multilayered bioinformatic approach that leverages genomic data across diverse species in combination with AI-based structural modeling to identify true ligand and receptor homologues, and subsequently predict their binding mechanisms. *In silico* findings were validated by multiple experimental approaches, which investigated the effect of amino acid changes in the proposed binding pockets on ligand-binding, complex formation with a co-receptor essential for downstream signaling, and activation of downstream signaling. Our analysis combining evolutionary insights, *in silico* modeling and functional validation provides a framework for structure-function analysis of other peptide-receptor pairs, which could be easily implemented by most laboratories.

## Introduction

Secreted signaling peptides are central regulators of growth, development, and stress responses in Eukaryotes. Plants, in particular, have evolved hundreds to thousands of such peptides and corresponding transmembrane receptors to regulate their growth and development in face of an ever-changing environment (1, 2). Knowledge of the exact binding mechanisms of these peptides to their transmembrane receptors is however limited to a handful of examples (3–11), mostly owing to the limited capability of most laboratories to perform structural analyses of ligand-receptor complexes. This ‘classical’ approach is indeed limited due to challenges with protein expression, purification and crystallization, and electron density maps can still be very difficult to obtain (12). Moreover, time, effort and cost of traditional crystal structure as well as cryo-EM determination approaches are of major constraint (13). Therefore, alternative approaches are urgently needed to study interactions at the receptor-ligand interface.

The family of stress-responsive SERINE RICH ENDOGENOUS PEPTIDES (SCOOPs) was identified in 2019 in the model plant *Arabidopsis thaliana* (hereafter, Arabidopsis) (14). Bioactive SCOOPs are 13-15 amino acid (AA) peptides proteolytically processed by subtilases from PROSCOOP precursors (14–17). Most Arabidopsis SCOOPs harbor the conserved ‘SxS’ motif that is essential for the bioactivity of the best characterized SCOOP, SCOOP12 (14, 15, 18). Recently, a comprehensive annotation of *PROSCOOP* genes in the Arabidopsis Col-0 genome revealed the existence of 50 putative SCOOP peptides, making the SCOOPs one of the largest families of signaling peptides identified in flowering plants so far (17).

Plants employ germline-encoded receptor kinases (RKs) and receptor proteins (RPs) to sense their extracellular environment and coordinate their growth and development in response to endogenous and exogenous cues (19). The most common ectodomain is a series of leucine-rich repeats (LRRs), which mediate ligand-binding and co-receptor association (20–23). SCOOPs were recently identified to be perceived by the LRR-RK MALE DISCOVERER 1-INTERACTING RECEPTOR LIKE KINASE 2 (MIK2) (15, 18).SCOOPs induce the complex formation between MIK2 and the common LRR-RK co-receptor BRASSINOSTEROID INSENSITIVE 1-ASSOCIATED KINASE 1 (BAK1) (15, 18). Notably, MIK2 and SCOOPs have been since implicated in multiple aspects of plant growth, development, and response to both biotic and abiotic stresses, therefore highlighting their biological relevance (14, 15, 17, 18, 24–30). Despite these advances, the evolutionary history of SCOOPs and MIK2, and how SCOOPs are perceived by MIK2 remain mostly unclear. The latter is particularly intriguing given that SCOOPs have divergent sequences apart from the conserved ‘SxS’ motif, and their sequences also suggest a different mode of binding compared to that of other characterized plant peptide-receptor pairs (20, 31, 32).

Multiple studies recently leveraged (pan)genomic data across and beyond plant families to gain insights into the structure-function mechanisms of RKs and RPs (33–37). Besides comparative genomics, protein structural modelling is now widely accessible. AlphaFold-Multimer (AFM) is an extension of AlphaFold2 (AF2) developed by DeepMind (38). Whereas AF2 predicts individual protein structures, AFM predicts structures of protein complexes with relatively high accuracy for ∼23% of the heteromeric interfaces (38). Although AF2 was only trained on monomer chains, it was quickly realized – owing to the idea that the molecular interactions governing protein folding are also of importance for protein-protein docking – that AF2 could also predict protein-protein models. Subsequently, AFM was released, an extension of AF2 specifically trained to predict protein complex structures with increased accuracy (12, 38, 39). Multiple *in silico* studies quickly reported AFM suitability for predicting peptide-protein interactions by challenging it against known interactors. Moreover, AFM outperforms state-of-the-art peptide-protein complex modeling (38, 40–45).

Here, we pioneer and functionally validate a relatively quick approach, combining the use of AFM and comparative genomics, to predict the ligand-binding pockets of an LRR-RK. Two binding pockets on MIK2 were predicted to interact with the conserved ‘SxS’ motif of the otherwise sequence-divergent SCOOPs identified within the plant order Brassicales. The AI predictions were supported by strong conservation of the predicted binding sites in novel validated MIK2 homologues across Brassicales. Site-directed mutagenesis of these binding pockets impaired SCOOP12-binding, SCOOP12-induced MIK2-BAK1 complex formation, and reactive oxygen species (ROS) production triggered by a multitude of SCOOPs.

## Results

### PROSCOOPs are exclusive to the plant order Brassicales, show diverse conservation patterns, and most harbor the ‘SxS’ motif

To shed light on PROSCOOP emergence and SCOOP conservation, a locus analysis was performed across 32 Brassicales species for each of the 19 Arabidopsis *PROSCOOP* loci that harbor the 50 Arabidopsis *PROSCOOPs* earlier identified (14–17). This strategy leveraged synteny and increased the number of putative *PROSCOOP*s to 381, facilitating a cluster analysis and the creation of 32 hidden Markov models (HMMs). The HMM profiles were then used as a query for an hmmsearch against 350 predicted proteomes across the entire plant kingdom and some unicellular algae (46). Manual curation of the resulting dataset settled on a total of 1097 putative *PROSCOOPs* identified in 32 species (Fig. 1A, Dataset S1).

**Fig. 1:**
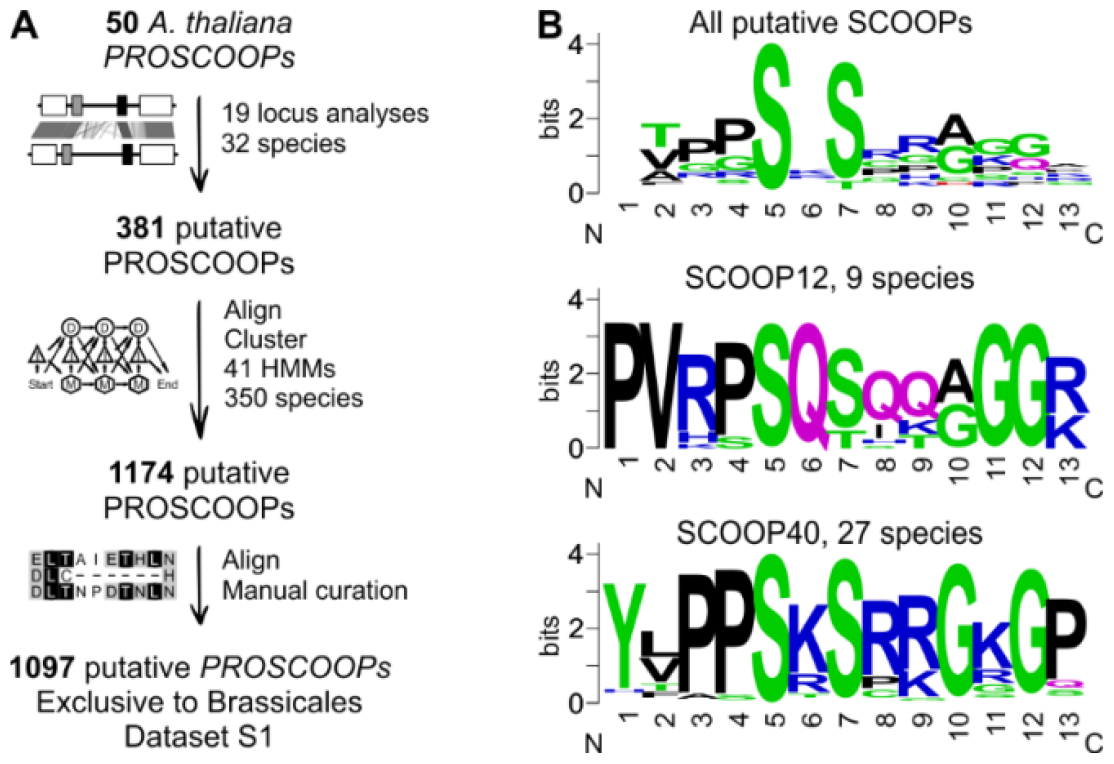
*In silico* mining of putative PROSCOOPs reveals that the majority harbors the ‘SxS’ motif, all PROSCOOPs are exclusively identified within the Brassicales. **A)** Schematic of the bioinformatic pipeline; a locus analysis facilitated PROSCOOP HMM-based mining across the plant kingdom, PROSCOOP candidates were subsequently manually curated. **B)** Sequence motif analysis of respectively all identified putative SCOOPs, SCOOP12 and SCOOP40. Sequence logos were generated using Dataset S1 and WebLogo server (https://weblogo.berkeley.edu/logo.cgi).

*Cleome violacea* is the earliest divergent species in which putative *PROSCOOP*s were identified. Three of them reside in the same clusters as Arabidopsis *PROSCOOP*40 and 48-49 (Dataset S1). Therefore, putative *PROSCOOP*s were exclusively found within Brassicales species, which diverged ∼39 mya (47). Five out of thirty-two maintained *PROSCOOP* clusters do not contain any Arabidopsis *PROSCOOP*s, suggesting that our bioinformatic pipeline was able to identify novel SCOOPs besides Arabidopsis SCOOP-homologues across species. Like the SCOOPs identified in Arabidopsis, Brassicales SCOOP sequences are divergent aside from the characterized ‘SxS’ motif, and a minority harbor an ‘SxT’ motif instead (Fig. 1B). Threonine (T), like serine (S), has a polar uncharged side chain and the ability to form hydrogen bounds. In addition, T also has a methyl group that allows the AA to establish van der Waals contacts with other non-polar groups..

In contrast to SCOOPs in general, motif analysis of SCOOP clusters reveals strong conservation of certain 13mer SCOOP sequences, suggestive of conserved function for some SCOOPs across species (Fig. 1B).

Although our search is biased by the annotation quality of the available genome assemblies (2), it allows us to acquire a general understanding of the sequence conservation of (PRO)SCOOPs within and across species. For example, clusters containing *PROSCOOP13* and *16, 37*-39, and *40* are represented in at least 27 out of 32 Brassicales proteomes, indicating a strong conservation after initial appearance during the evolution of Brassicaceae. In contrast, *PROSCOOP5, 34* and *43* sequences seem to be relatively unique as they did not cluster with any other sequences post the initial locus analysis within 32 species. *PROSCOOP29, 30, 42, 44, 45, 46* and *47* clustered with just one other sequence. *PROSCOOP2, 7, 12* and *14* are found in less than 10 out of 32 Brassicales species. Independent of their evolutionary conservation within the Brassicales, 13mer SCOOP sequences can be strongly conserved (Dataset S1).

To test the conservation of plant responses to SCOOPs, we measured the production of reactive oxygen species (ROS) – a hallmark of LRR-RK activation – triggered by SCOOPs. SCOOP12 was used as it is the best characterized SCOOP (14, 15, 18, 25, 26). Moreover, Arabidopsis SCOOP12 was earlier shown to induce apoplastic ROS production in *Brassica napus* (14), even though our dataset indicates that *B. napus* does not have a *bona fide PROSCOOP12*-homologue in the cluster containing Arabidopsis PROSCOOP12. Moreover, relative to SCOOP12, the maximum AA similarity with any predicted *B. napus* SCOOP (13 AA) is∼54 % (SCOOP27 cluster, Dataset S1). Besides SCOOP12, SCOOP13, 16, and 24 were selected for testing, based on the following three criteria: A) relatively conserved across the Brassicales, B) strong sequence conservation of the predicted active SCOOP, and C) induction of robust ROS production upon SCOOP treatment in Arabidopsis (17). Brassicaceae plants were carefully selected to cover the diversity of the plant family (Fig. S1). *C. violacea* was selected as a close outgroup for the family of the Brassicaceae (order of the Brassicales). We measured ROS production following application of the selected SCOOPs in these plants and included the immune elicitor flg22 as a positive control to test the suitability of our assay for each species. *Carica papaya*, a species from a relatively close outgroup for the Brassicales, did not respond to flg22 in our ROS production assay and was therefore excluded from further analyses. Besides Arabidopsis, *Brassica rapa, Eutrema salsugineum, Euclidum syriacum, Diptychocarpus strictus* and *C. violacea* responded significantly to at least two out of four tested SCOOPs (Fig. S2). Hence, phylogenetic and experimental evidence suggests that SCOOP perception occurs within the Brassicales.

### MIK2 is a Brassicales-specific, conserved SCOOP receptor

To explore the gain of SCOOP response in relation to its defined receptor in Arabidopsis, putative MIK2 homologues were identified *in silico* following a multilayered approach similar to the (PRO)SCOOP mining. Initially, leveraging synteny, a locus analysis identified 38 putative MIK2s within 32 Brassicales species. The advantage of locus analysis against genome-wide searches is the probable common evolutionary origin of the gene of interest. Subsequently, an HMM profile was created and used to interrogate 350 species across the plant kingdom (46). In this way, we also included putative MIK2 homologues and putative MIK2 paralogues RKs within and outside the Brassicales, which do not necessarily reside within the conserved *MIK2* locus. Subsequently, a phylogenetic analysis was performed and a clade containing all initial putative MIK2s was extracted. After manual curation of the alignment (24 LRRs, no gaps in other conserved domains), 17 novel LRR-RKs were identified besides the earlier identified 38. As a relatively close outgroup of the Brassicales, we also investigated the *MIK2* locus of *Carica papaya* and found that the *MIK2* locus of *C. papaya* does not contain any LRR-RK-encoding gene (Fig. 2A). In contrast, earlier diverged species such as *Theobroma cacao* do harbor one or more LRR-RK-encoding genes at the *MIK2* locus, but with a relatively low sequence similarity to Arabidopsis MIK2 (Thecc08G107800: 61 %). In summary, no putative MIK2 homologues were identified outside the Brassicales.

**Fig. 2:**
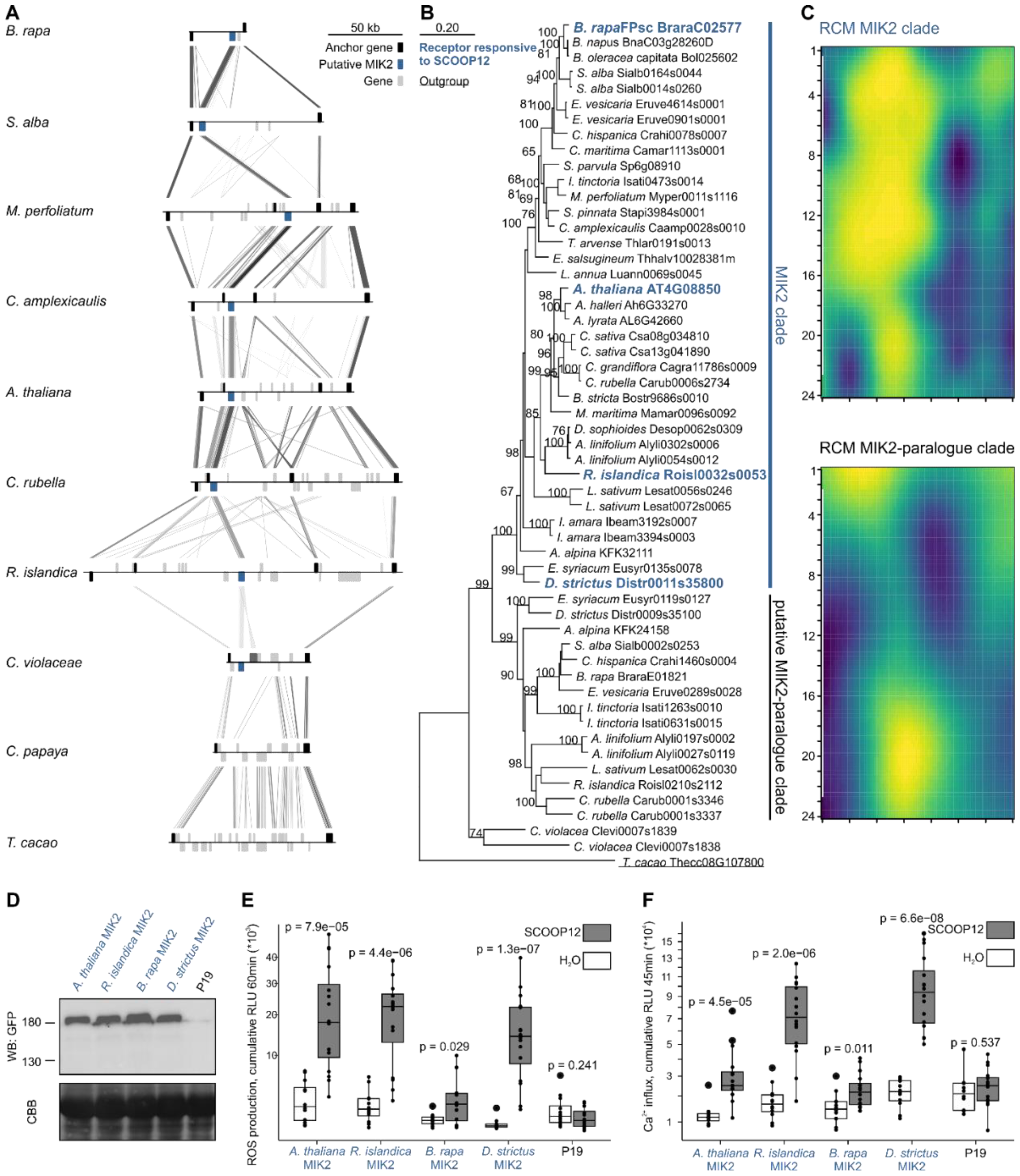
MIK2 is specific to and strongly conserved within the Brassicales family. **A)** Anchor, putative *MIK2*, and other genes are colored as per legend. Locus comparison of eight contiguous MIK2 loci within the Brassicales and the relatively close outgroups *C. papaya* and *T. cacao*. Blast hits between loci are indicated with lines (e-value <1e-06) with score according to grayscale gradient and darker grays indicating higher similarity. Gene orientation is depicted by the position of them above or below the line. **B)** A phylogenetic analysis of putative MIK2 homologues and paralogues. Maximum likelihood analysis bootstrap values are indicated, and only values higher than 65 are shown. The scale bar represents 0.2 AA substitutions per site. An LRR-RK that resides at the conserved MIK2 locus of *T. cacao* was used as an outgroup to root the phylogenetic gene tree and is underlined. The functionally validated MIK2 receptors of *A. thaliana, R. islandica, B. rapa* and *D. strictus* are highlighted in green and bold as they confer SCOOP-induced reactive oxygen species (ROS) production and increase in cytosolic Ca^2+^ concentration upon heterologous expression in *N. benthamiana*, as shown respectively in panel E and F and Fig. S3. These four validated MIK2s fall within the labeled ‘MIK2 clade’. **C)** Repeat conservation mapping (RCM) of the putative MIK2 (n=37) and potential MIK2 paralogues (n=15) clade. Each row represents the solvent exposed AAs of a single repeat of the LRR. The color represents the center-weighted regional conservation score for the 5x5 set of boxes that centers on that box; yellow indicates the most conserved regions and blue indicates the most divergent regions. **D)** Western blot of the heterologously expressed MIK2 and MIK2 homologues in *N. benthamiana*. Tissue was harvested 48 h after construct infiltration in *N. benthamiana*. The western blot was probed with α-GFP (B-2) HRP as the receptor had a C-terminal GFP tag (top) and subsequently stained with CBB as a loading control (bottom). See Dataset S6 for original tiff files. **E-F)** Shown are ROS production (E) and increase in Ca^2+^ cytosolic concentrations (F), respectively 4-60 min and 3-45 min, in cumulative relative luminescence units (RLUs) post treatment with H_2_O (white) or the peptide SCOOP12 (1 μM, grey). Four independent biological replicates (n=4 plants) were performed, with each biological replicate represented by four technical replicates. Significance was tested by performing a paired Wilcoxon rank-sum test.

Two maximum-likelihood phylogenetic analyses – using either sequences of the extracellular domain only or the full protein – were performed with the 54 putative MIK2s. Thecc08G107800 (from *T. cacao*) was included in the analysis to root the tree (Fig. 2B). Both strategies identified a putative MIK2 clade containing 37 putative Brassicaceae MIK2 homologues, all residing at the *MIK2* loci, with an AA similarity of 86-97 % relative to Arabidopsis MIK2. A common function of all LRR-RKs within the putative MIK2 clade is likely as a Repeat Conservation Mapping (RCM) analysis (48) identified conserved sites in a putative ligand-binding region spanning LRR1-15 on the predicted surface of the LRR ectodomain (Fig. 2C) (35).

In contrast, 16 out of 17 novel MIK2 candidates cluster in a distinct clade, show a relatively low conservation in the RCM between LRR1-15 (Fig. 2C), and none of these LRR-RK-encoding genes reside at one of the earlier identified *MIK2* loci. All members within this clade, hereafter described as the putative MIK2-paralogue clade also have a relatively lower AA similarity with Arabidopsis MIK2 of 74-81 %. The last novel MIK2 candidate, Clevi0007s1838, resides at the *MIK2* locus of *C. violacea* (Brassicales) and clusters together with its neighboring gene *Clevi0007s1839* but separate from all other putative MIK2-homologues and -paralogues within the Brassicaceae. Clevi0007s1839 shares approximately 79 % AA sequence similarity with Arabidopsis MIK2.

Next, we investigated the function and relationship of individual putative MIK2 homologues within the putative MIK2 clade. Besides Arabidopsis *MIK2*, three genes – *Roisl0032s0053* (from *Rorippa islandica*), *BraraC02577* (from *B. rapa*) and *Distr0011s35800* (from *D. strictus*) – were selected as representatives for the major subdivisions within the Brassicaceae, cloned, and transiently expressed in the non-Brassicaceae model species *Nicotiana benthamiana*, which is insensitive to SCOOPs (14, 18). As in previous experiments testing Brassicales species for SCOOP responsiveness, we initially used SCOOP12 as it is the best characterized SCOOP to validate the function of our putative MIK2s and the putative MIK2 clade in general. Receptor function was measured using SCOOP-induced ROS production and increase in cytosolic Ca^2+^ concentration. Transient expression of all three representative genes conferred SCOOP12-induced activation of both tested hallmarks of LRR-RK signaling, consistent with SCOOP recognition enabled by the MIK2-clade (Fig. 2 E,F). Similar to our analysis of SCOOP perception across species, we then tested whether transient expression of the three MIK2 homologues could confer response to additional SCOOPs, namely SCOOP13, 16 and 24. Arabidopsis SCOOP13 and 16 differ by two AAs, are part of the same cluster that was found in 28 species and have a strong sequence conservation across them. Similar ROS bursts were observed for both SCOOPs as all MIK2 homologues conferred responsiveness except *B. rapa* MIK2 (BraraC02577) (Fig. S3). Similarly, transient expression of *B. rapa* MIK2 did not lead to significant ROS production [Wilcoxon-ranked sum (WRS), p=0.14] upon treatment with SCOOP24, a strongly sequence-conserved SCOOP found in 22 Brassicaceae (Dataset S1). Nevertheless, all tested SCOOPs did result in an increase of cytosolic Ca^2+^ concentration, often a more sensitive marker for RK signaling relative to ROS production. In summary, SCOOP perception across the Brassicales is correlated with the *in vivo* function of MIK2 homologues.

The conserved SCOOP motif ‘SxS’ is predicted to interact with two MIK2 binding pockets To understand how MIK2 perceives SCOOP ligands, we used AFM in combination with RCM. Because peptide ligands are perceived by LRR ectodomains, and since AF2 does not correctly orient ecto- and cytosolic domains with respect to transmembrane domains (49), we elected to predict the interaction between SCOOPs and the isolated MIK2 ectodomain. We did not include the co-receptor BAK1 as it only gets recruited in the SCOOP12-MIK2 complex upon ligand binding (15, 18). Building on our evolutionary analysis of MIK2 and SCOOPs, we hypothesized that the SCOOP ‘SxS’ motif forms contact with the putative ligand-binding site predicted in our RCM analysis (Fig. 2C). AFM predicts a high interface predicted Template Modelling (ipTM>0.84) score for twelve out of fifty Arabidopsis SCOOPs in complex with Arabidopsis MIK2 (Fig. 3A). Strikingly, the predicted interactions between SCOOP12 and MIK2 all fall within the strongly conserved putative functional sites for ligand recognition on the predicted surface of the LRRs, as delineated in our RCM analysis (Fig. 3B). Across species and SCOOPs, only the ‘SxS’ motif is relatively strongly conserved. Moreover, single serine to alanine mutations in the ‘SxS’ motif of SCOOP12 either reduce or abolish SCOOP12-induced ROS in Arabidopsis (14). In the AFM model, the ‘SxS’ SCOOP motif is predicted to interact with the same two putative binding pockets on MIK2 across all twelve high-scoring AFM predictions. The hereafter referred to as S5 and S7 MIK2-binding pockets engage in hydrogen bond interactions with S5 and S7 of the 13mer SCOOP peptide, and play a key role as recognition points for the peptide within the receptor. These pockets are composed by the MIK2-specific AAs D246/N268 and S292/H294/ H316, respectively (Fig. 3C). Consistent with the AFM complex predictions, these two binding pockets are fully conserved within the earlier defined MIK2-clade as earlier suggested by the RCM. We additionally attempted to predict a receptor/co-receptor/ligand complex for the 12 SCOOPs that resulted in a high ipTM with solely the receptor, again using just the ectodomains, but AFM failed to predict the tripartite MIK2-BAK1-SCOOP complex without any AA side chain clashes.

**Fig. 3:**
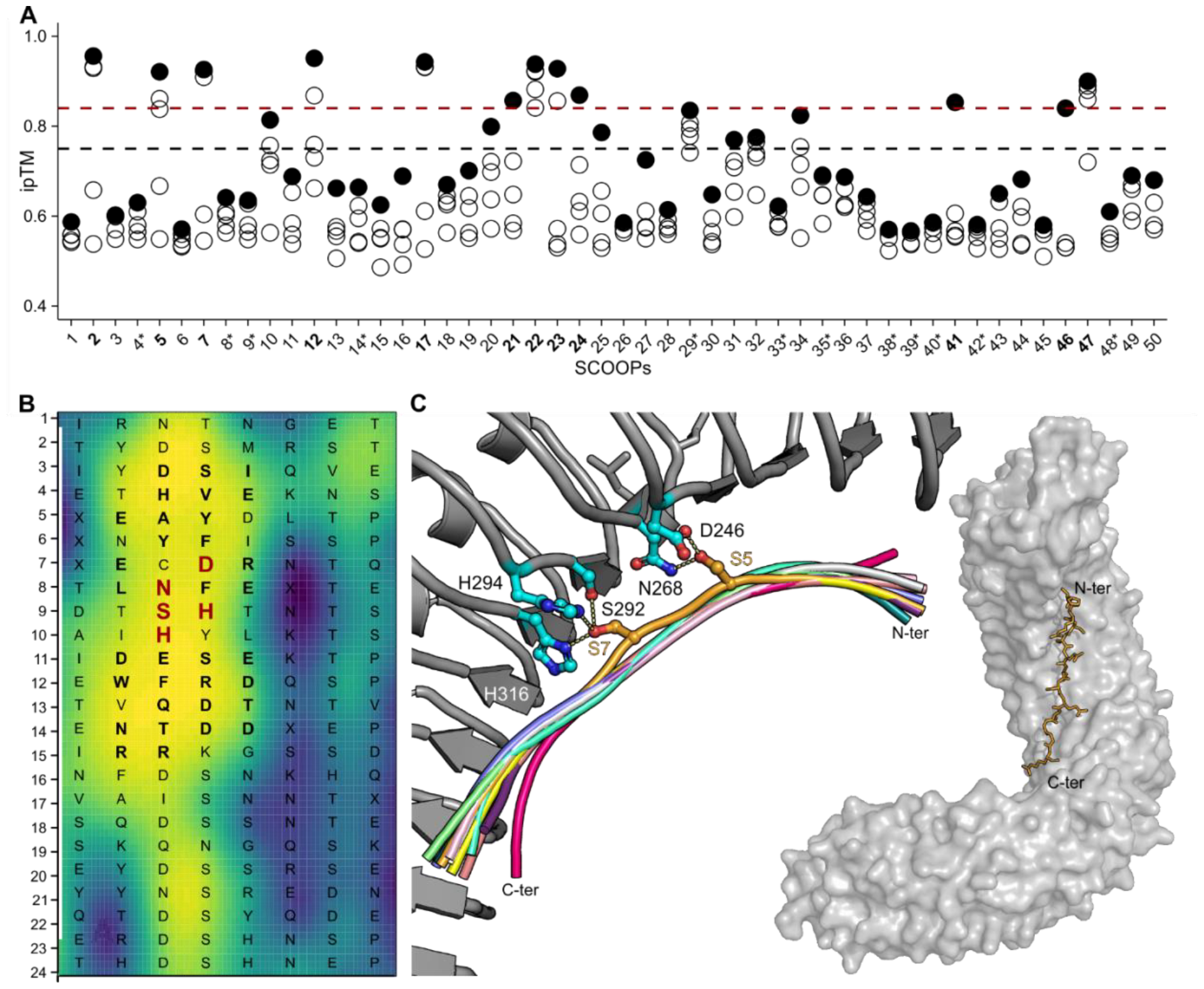
Alphafold-multimer (AFM) predicts that the ‘SxS’ motif interacts with two conserved binding pockets of MIK2. **A)** Alphafold-Multimer (AFM) predicts a high interface predicted Template Modelling (ipTM) score for 12 (bold) out of 50 Arabidopsis SCOOPs in complex with the MIK2 receptor. Asterisks indicate the use of the 15mer instead of 13mer SCOOP for complex prediction. **B)** Repeat conservation mapping (RCM) of the putative MIK2 (n=37) clade. Each row represents a single repeat of the LRR, with each colored box representing a solvent exposed amino acid position. The consensus sequence of the 37 MIK2s is depicted, single AA are enlarged and in bold in case an interaction with SCOOP12 was predicted by AFM, and additionally highlighted in red in case AFM predicts an interaction with the conserved ‘SxS’ motif of SCOOPs. The color represents the center-weighted regional conservation score for the 5x5 set of boxes that centers on that box; yellow indicates the most conserved regions and blue indicates the most divergent regions. **C)** Structural superposition of 11 SCOOPS in cartoon representation (12, 2, 5, 7, 17, 21, 22, 23, 24, 46, 47) with the 2 conserved S (depicted in orange sticks) anchoring the peptide to the receptor binding canyon. S5 mediates hydrogen bond interactions with the conserved residues D246 and N268 (depicted in cyan sticks). S7 conserved binding pocket is composed by S292, H316 and H294 (depicted in cyan sticks). On the right side, an AFM model prediction of MIK2 in complex with SCOOP12 is depicted. SCOOP12 is predicted to bind to the MIK2 internal binding groove in a fully extended conformation. MIK2 is depicted in grey surface and SCOOP12 in yellow sticks.

### Single AA changes within the AFM-predicted binding pockets affect ligand-induced ROS production

To test experimentally the biological relevance of the predicted binding pockets, we created constructs with single AA substitutions in both the S5 and the S7 binding pockets. We expressed wild-type Arabidopsis MIK2 or binding-pocket variants in *N. benthamiana*, and tested the capacity of the variants to perceive SCOOPs as measured by SCOOP-induced ROS production. Western blot analysis and confocal microscopy demonstrated that most variants accumulated to comparable protein levels with wild-type MIK2 and were correctly localized to the plasma membrane, respectively (Fig. S4). One variant, H316G, exhibited altered subcellular localization and was therefore excluded from further experiments. The SCOOPs tested were selected from diverse Arabidopsis SCOOP clusters and are known to induce ROS production in Arabidopsis (17).

Intriguingly, single AA changes in the S5 binding pocket consistently reduced or abolished MIK2 function (Fig. 4A and Fig. S7). The D246G MIK2 variant abolished ROS production in response to all tested SCOOPs, whereas N268G abolished ROS in response to 7 out of 8 tested SCOOPs and reduced it in response to SCOOP5 relative to WT MIK2 (WRS, p=0.0059). S292G from the second binding pocket abolished ROS production in response to 6 out of 8 SCOOPs and reduced it for SCOOP18 and 21 (WRS, respectively p=0.0168 and p=3.364e-05). This indicates the occurrence of at least a certain level of ligand binding of the respective SCOOPs followed by complex formation with BAK1. H294G abolished ROS production in response to only 4 out of 8 tested SCOOPs, did not result in a significant difference for SCOOP5, 12 and 18 (WRS, respectively p=0.0972, p=0.9852 and p=0.2099), and unexpectedly resulted in stronger ROS production in response SCOOP24 relative to WT MIK2 (WRS, p=3.412e-05).

**Fig. 4:**
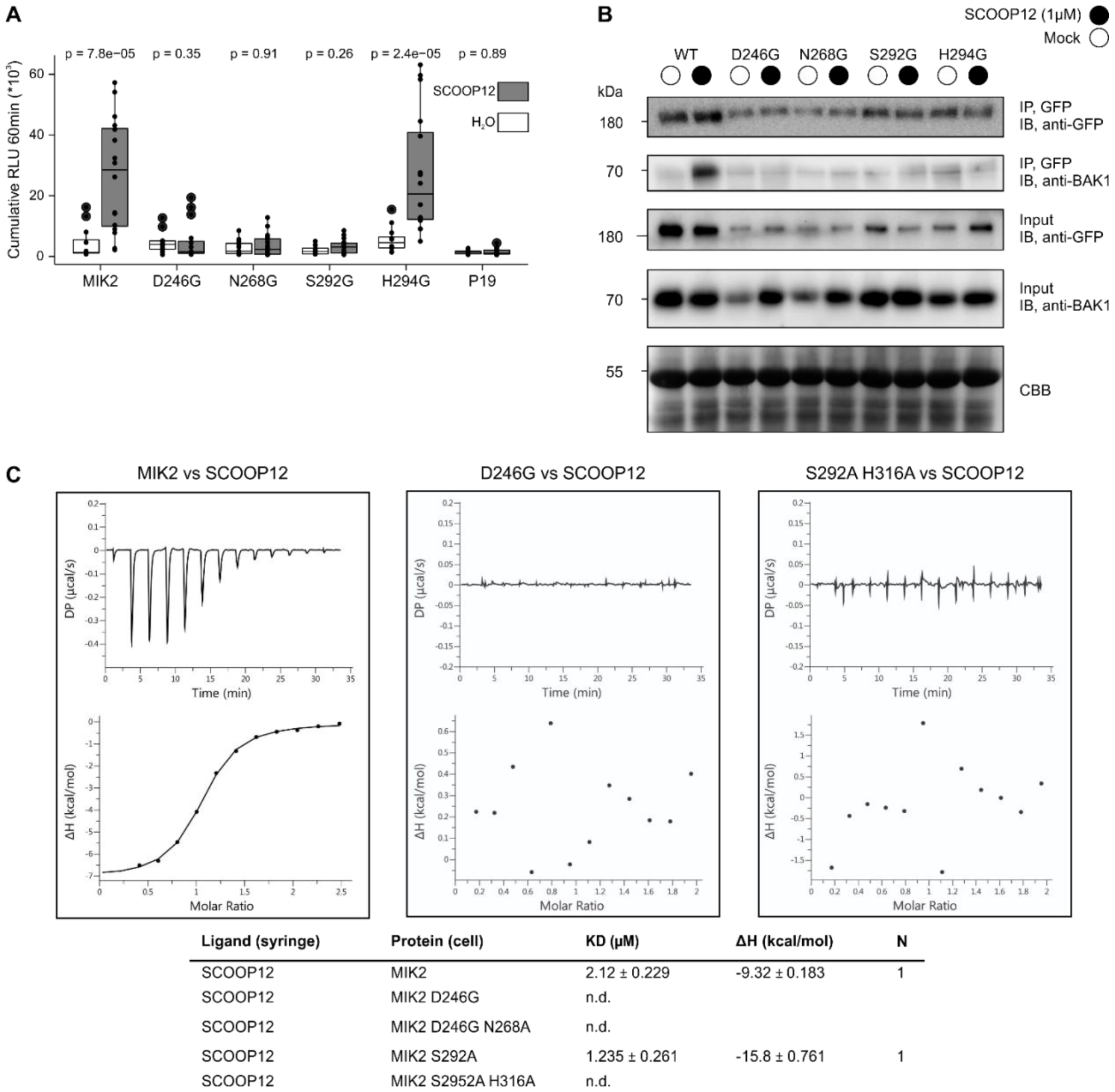
Single and double AA changes within the putative MIK2 binding pockets affect SCOOP12 ligand binding, SCOOP12-induced MIK2-BAK1 complex formation and ROS. **A)** Shown is ROS production (4-60 min) in cumulative relative luminescence units (RLUs) post treatment with H_2_O (white) or SCOOP12 (1 μM, grey). Four independent biological replicates (n=4 plants) were performed, with each biological replicate represented by four technical replicates. Significance was tested by performing a paired Wilcoxon rank-sum test. **B)** Co-immunoprecipitation post heterologous expression in *N. benthamiana* of BAK1 with MIK2-GFP after treatment with 1 μM SCOOP12, or water for 15 min. Western blots were probed with antibodies α-GFP and α-BAK1. **C)** Isothermal titration calorimetry (ITC) experiments of MIK2 binding pockets variants vs SCOOP12 and summary table. *K*_d_ (dissociation constant) indicates the binding affinity between the two molecules considered (in micromolar). The N indicates the reaction stoichiometry (*n* = 1 for a 1:1 interaction). The values indicated in the table are the mean ± SD of at least two independent experiments. N.d. = non detected binding.

### Single AA changes within the AFM-predicted binding pockets affect MIK2-BAK1 complex formation

Like many other LRR-RKs, ligand binding to MIK2 triggers complex formation with co-receptor kinases from the SOMATIC-EMBRYOGENESIS RECEPTOR-LIKE KINASE (SERK) family, primarily BAK1 (15, 17, 18). A co-immunoprecipitation assay was performed after heterologous expression in *N. benthamiana* to test whether MIK2 variants could form a complex with BAK1 following SCOOP perception (Fig. 4B). Relative to the corresponding mock treatments, a clear induction of complex formation could be observed for MIK2-BAK1 post SCOOP12 treatment. This was not the case for MIK2 variants. However, it is important to highlight that this is a semi-quantitative assay, and we cannot exclude that complex formation for MIK2 variants happens at a lower level relative to WT MIK2.

### Single AA changes within the AFM-predicted binding pockets affect ligand binding

To investigate the relevance of the S5 and S7 MIK2 binding pockets in the anchoring and recognition of SCOOP12 to the receptor; we produced recombinant ectodomains of MIK2 binding variants in both pockets, using insect cells cultures. These variants were then subjected to direct isothermal titration calorimetry (ITC) experiments with synthetic SCOOP12. We designed and tested the following mutants in the distinct MIK2 binding pockets: D246G/N268A and D246G in the S5 pocket and S292A and S292A/H316A in the S7 pocket. Due to the incorrect localization observed in the full-length H316G MIK2 mutant in the S7 binding pocket in *N. benthamiana* (Fig. S4), we generated a new variant where H316 was substituted with alanine (A). All expressed MIK2 mutants exhibited proper folding and eluted as monomers in size-exclusion chromatography experiments (Fig. S5). To evaluate the interaction between MIK2 pocket variants and SCOOP12, we titrated the peptide into a solution containing the isolated MIK2 ectodomain variants. We did not detect any binding of SCOOP12 to the double MIK2 mutant D246G/N268A in the S5 binding pocket (Fig. 4C and Fig. S6). We next assessed the individual contribution of the core S5 binding pocket residue D246. ITC experiments revealed that mutation of D246 alone is sufficient to disrupt the anchoring and recognition of SCOOP12 by MIK2 (Fig. 4C and Fig S6). In contrast, the single mutation S292A in the S7 pocket retained the ability to bind the peptide with WT affinity (Fig. 4C, (18)). However, when the S292A mutation was combined with H316A, the interaction with SCOOP12 was completely lost (Fig. 4C and Fig S6). These *in vitro* biochemical data therefore confirm our computational prediction as well as validated *in vivo* biochemical and physiological data.

## Discussion

Secreted signaling peptides regulate growth, development, and stress responses. In this study, we described the evolution of a lineage-specific phytocytokine family and its receptor. Subsequently, we leveraged the acquired receptor/peptide homologues in combination with AI-driven protein structural prediction to unravel the mechanism of ligand binding. Our approach paves the way for rapid identification of peptide-receptor interaction mechanisms.

Initially, *PROSCOOP12* was identified in Arabidopsis by analysis of transcriptomic profiles upon exposure to stresses (14). A subsequent screening of the genome revealed a novel peptide family that resides on just two loci that harbor 14 homologous genes with a similar intron-exon structure. Seven additional species were mined for putative homologues using a BLASTP approach, resulting in 74 putative *PROSCOOPs* within the Brassicaceae (14). The list of Arabidopsis *PROSCOOPs* was then extended to 23 and then 28 members (15, 16). However, a recent comprehensive bioinformatic analysis brought the total number to 50 diverse *PROSCOOPs* (17). It is generally difficult to identify peptide homologues from distantly related species by BLAST, due to the high sequence variability of prepropeptides, except the short region encoding the mature peptide. Hence, it is advised to limit the query to the most conserved part of the prepropeptide (2). Lastly, due to the low similarity between putative homologues, it is unclear whether they are true homologues, or just sequences that evolved independently (50). Therefore, this study leveraged a locus analysis (∼evolutionary linkage), which facilitated subsequent HMM searches with optimized queries across 350 predicted proteomes covering the plant kingdom. This effort resulted in 1097 putative *PROSCOOPs* (Dataset S1), transcending the Brassicaceae family, but limited to the order of the Brassicales. Hence, the SCOOP family, like it is proposed for the systemin family, is evolutionary young relative to three other well-characterized stress-related secreted signaling peptide families, PIPs, PEPs and CTNIPs/SCREWs, which were identified across a multitude of diverse Angiosperms (2, 51–55). However, in contrast to systemin (55), the SCOOP family is ubiquitously present within the lineage they occur.

Most putative Brassicales SCOOPs have the previously described ‘SxS’ motif (Fig. 1), and serine to alanine mutations in SCOOP12 highlighted the importance of these two serine residues for SCOOP perception (14). The double S5A/S7A, and single S5A mutation did not induce ROS production whereas the S7A mutation resulted in a low, but still significant ROS production. Moreover, ITC analysis showed that the MIK2 ectodomain binds SCOOP12 but not the double S5A/S7A SCOOP12 variant (15). The SCOOP ‘SxS’ motif is unique across known plant secreted signaling peptides in contrast to the N-terminal asparagine and sulfated tyrosine motif found in RGF, PSY and CIF peptides and the core PSGP sequence of the proline-rich CLE, CTNIP, PIP, PIPL, CEP and IDA+IDL peptides (56). In contrast to the broader SCOOP family, certain individual SCOOPs show a strong conservation across the length of the predicted active peptide (Fig. 1B, Fig S2B), suggesting a conserved function across the species in which they were identified.

Although evolutionary analysis of secreted signaling peptide receptors has been reported previously (51, 52), earlier studies lacked the depth required to study the emergence of specific receptor functions and facilitate their mechanistic understanding (33, 57). Moreover, with more genomes sequenced, there is presently great opportunity to explore peptide signaling beyond model species (56). We measured SCOOP-induced ROS production across the order of the Brassicales and observed it in all species tested. In contrast, *N. benthamiana* and *Solanum lycopersicum* from the Solanaceae family are non-responsive to SCOOP12 treatment (14).

We complemented phenotypic observations with a multitude of *in silico* approaches to unravel the evolutionary gain of MIK2 and SCOOP perception. Analysis of the *MIK2* locus across 32 species facilitated an HMM search across 350 predicted proteomes. Phylogenetic analysis using the kinase domains of the resulting LRR-RKs delineated a monophyletic clade, which contained all previously identified putative MIK2 homologues. After manual curation – as it is important to filter for an equal amount of LRRs when identifying LRR receptor homologues (2) – a maximum-likelihood phylogeny was performed, which ultimately revealed a putative MIK2 clade containing 37 LRR-RKs from 31 species including *A. thaliana* MIK2. The putative MIK2-paralogue clade is the closest related clade to the putative MIK2 clade, contains 15 putative MIK2 paralogues – all residing outside the contiguous MIK2 locus – and are only found in species that also contain a putative MIK2 homologue. Finally, we performed a RCM analysis on both clades of putative MIK2 homologues and paralogues (48). RCM predicts functional sites in LRR domains using signatures of conservation/diversification of surface residues, given a group of receptor homologues as an input. Hence, the presence of shared predicted functional sites within putative receptor homologues such as the putative MIK2 clade is an indicator that they might share a conserved function.

LRR-RKs of the putative MIK2 clade of four Brassicaceae genera were tested for SCOOP responsiveness and could induce ROS production and elevated cytosolic Ca^2+^ concentrations upon heterologous expression in *N. benthamiana*. Intriguingly, none of the investigated Brassicaceae lacks a MIK2 homologue within the MIK2 clade, suggestive of a strong conservation subsequent to the evolutionary gain of SCOOP perception. Neither MIK2 homologues, nor putative MIK2 paralogues, were identified outside the Brassicales. Hence, combining the results of the *in silico* analysis of SCOOPs and MIK2, the native plant responses of Brassicales, and the response of diverse MIK2 homologues post heterologous expression upon treatment with SCOOPs, we suggest the appearance of ancestral SCOOPs and an ancestral MIK2 at least ∼39 mya (47). However, more genome assemblies of species that diverged relatively closely to the divergence of the Brassicales species *C. violacea* are crucial to resolve the exact sequence of these events.

Additionally, our functional analysis of MIK2 homologues in a heterologous model (*N. benthamiana*) provides insight into LRR-RK function as it facilitated an in-depth RCM analysis. Additional sequences increase the reliability and power of RCM analysis. Hence, using 37 MIK2 homologues, this resulted in a clear distinction between conserved and diversified areas. Conserved regions on the surface of folded proteins often correspond to key functional sites such as for example ligand-binding sites (48, 58, 59). Nevertheless, not all interacting residues of LRR receptors and ligands are necessarily conserved across homologues as they might confer specificity. For example, although present across species, neither tested CTNIPs nor PEPs are recognized across species boundaries (a phenomenon referred to as con-specificity), putatively due to co-evolution of the ligand and its receptor (51, 52). Additionally, not all high-scoring RCM residues interact with ligands based on the available structural data of receptor-ligand complexes (48). Lastly, interpretation of RCM analysis is easier with the help of a protein structural model. Therefore, we opted to combine this strategy with a novel approach, the use of AI-driven protein structural prediction with AFM. The conserved SCOOP ‘SxS’ motif was predicted to interact with two conserved binding pockets within all MIK2 homologues. Moreover, MIK2 residues predicted to interact with the 13mer SCOOP12 all fall within conserved RCM residues (Fig. 3B). Hence, these two diverse approaches strengthen each other’s predictions as they point towards the same putative binding area.

The S5 and S7 binding pockets were functionally validated by testing MIK2 variants using three experimental approaches; A) SCOOP12-binding to MIK2 ectodomain *in vitro*, B) SCOOP12-induced complex formation with its co-receptor BAK1 *in vivo*, and 3) ROS production upon SCOOP treatments as a marker for receptor complex activation (Fig. 4, Fig. S7). For example, relative to WT MIK2, a single AA mutation of D246 within the S5 binding pocket abolished direct SCOOP12 binding, diminished complex formation with BAK1 upon SCOOP12 exposure and abolished ROS production upon heterologous expression in response to eight diverse SCOOPs. In contrast, the S292A AA mutation within the S7 binding pocket did not abolish SCOOP12 binding *in vitro* but the double mutation S292A H316A did. Moreover, the two single mutations in the S7 binding pocket have less drastic impact on ROS production to certain SCOOPs. This suggests that complex formation with BAK1 might still happen depending on the interacting SCOOP and the specific mutation (for example H294G and SCOOP12), although at a relatively low but sufficient level as indicated by the co-immunoprecipitation assay. Importantly, these results are consistent with single replacements to alanine within the ‘SxS’ motif of SCOOP12. Whereas SCOOP12 S5A does not induce ROS production in Arabidopsis Col-0, SCOOP12 S7A shows a comparatively low but significant ROS burst (14). Finally, although most putative SCOOPs contain the ‘SxS’ motif, a minority harbor an ‘SxT’ motif instead (Fig. 1). Therefore, we hypothesize that the S5 binding pocket functions as an anchor point by initiating SCOOP binding through S5 of the ‘SxS’ motif. Subsequently, the S7 binding pocket most likely stabilizes SCOOP binding. Intriguingly, in contrast to other peptide families that can be perceived by several phylogenetically related LRR-RKs, perception of the 50 predicted SCOOPs seems to solely necessitate MIK2 (17, 18). Differentially affected ROS production by MIK2 variants following exposure to certain SCOOPs suggest that SCOOP-induced responses might partly rely on divergent binding of SCOOPs to MIK2 besides potential transcriptional and spatial regulation (60).

In this study, beyond deciphering SCOOP/MIK2 co-evolution and SCOOP-MIK2 binding mechanisms, we pioneered the use of AI-driven protein complex prediction by AFM in combination with comparative genomics to identify ligand-binding pockets for a peptide-receptor pair. The success of our approach depends on the accuracy of the complex prediction, which remains a challenge relative to monomer predictions (43). Not surprisingly, using the standard AFM approach, the interaction of the ‘SxS’ motif of 12 out of 50 SCOOPs with MIK2 were predicted correctly. Nevertheless, results of the 5^th^ joint CASP-CAPRI protein assembly prediction challenge indicate a remarkable improvement of complex predictions relative to the 4^th^ meeting two years prior (43). Moreover, a multitude of participating groups exceeded the performance of the benchmark standard, AFM. Hence, AI-driven complex predictions will certainly improve and thus will play an important role in unravelling other peptide-receptor interactions in the future.

## Material and Methods

### PROSCOOPs

#### Mining and analysis

First, a locus analysis was performed for each Arabidopsis *PROSCOOP* locus across the Brassicales as described before (17, 57). The analyzed genome assemblies, versions, and their sources can be found in Dataset S2. In short, BLASTP (BLAST 2.9.0+, e-value 10) was used to identify *PROSCOOP* syntenic loci by mining the genomes for homologues of the strongly conserved neighbor genes of Arabidopsis *PROSCOOPs* which can be found in Dataset S2. Second, the novel candidate PROSCOOPs were added to the earlier identified 50 Arabidopsis PROSCOOPs and clustered with MMseqs2 (release 14-7e284) using a minimal sequence identity and coverage of 0.5 and 0.3, respectively (61). Third, for each cluster with at least three sequences, we built an HMM profile by aligning the sequences with muscle and running hmmbuild (62, 63). These profiles were used to search for additional *PROSCOOP* candidates in a collection of genomes from 350 species described previously (Ngou et al., 2022). The results were filtered for an e-value below 10e-5, resulting in a total of 1168 *PROSCOOP* candidates, all are limited to Brassicales. Finally, putative SCOOPs were extracted based on the corresponding 13- and 15mer of the Arabidopsis SCOOPs if present in the cluster, otherwise the position of the ‘SxS’ motif was used to predict a 13mer. Putative active SCOOPs were extracted for 1097 out of 1168 sequences. Seventy-one candidate *PROSCOOP*s were filtered out due to irregularities in the alignment potentially caused by ORF shifts due to sequencing errors, incorrect annotations, or pseudogenes. Gene accessions, protein sequences and predicted 13-mers of all putative *PROSCOOPs* can be found in Dataset S1.

### MIK2

#### Mining and analysis

Similar to the *SCOOP*s, the analyzed genome assemblies, versions, and their sources included in the contiguous *MIK2* locus analysis can be found in Dataset S2. Putative MIK2 homologues were identified using locus analysis within 32 Brassicales species. In short, BLASTP (BLAST 2.9.0+, e-value 10) was used to identify the *MIK2* syntenic locus and putative MIK2s by mining the genomes initially for homologues of the strongly conserved neighbor (anchor) genes of Arabidopsis *MIK2* (*AT4G08850*); *AT4G08810*/*AT4G08840* and *AT4G08870*/*AT4G08920*. Subsequently, putative MIK2 homologues were identified using BLASTP, resulting in a list of 39 putative MIK2 homologues. Locus comparison was performed using R (v4.0.3) and the R-package genoPlotR (v0.8.11) using the extracted contiguous *MIK2* loci and their corresponding annotation (Dataset S2). The resulting figure was edited in Corel-DRAW Home & Student x7.

Subsequently, we built an HMM profile with hmmbuild (version 3.1b2) and searched all proteins from the collection of 350 genomes that were longer than 300 AA for matches (hmmsearch --max -E1e-10) (63). Given the relaxed e-Value threshold, we found 435,387 initial matches which is close to what one would find with a plain protein-kinase HMM from PFAM. We then filtered for matches with an e-value below 1e-250, thus reducing the set to 4,786 candidates. All initial MIK2 homologues identified through the synteny approach were included in the novel list. We then extracted the kinase domain using hmmsearch and the kinase PFAM pattern PF00069.26. For each sequence, we selected the best matching stretch and extracted the sequence with bedtools (version 2.92.2) (64). Sequences were then aligned with FAMSA (version 1.6.2) (65). Alignments were not trimmed, and phylogenetic trees were inferred with FastTree (version 2.1.11 SSE3, option -lg) (66, 67). Finally, we extracted the clade that contained all initial candidates with gotree (version 0.4.0, github.com/evolbioinfo/gotree).

#### Maximum-likelihood Phylogeny

The above set of putative *MIK2* homologues and potential *MIK2* paralogues was filtered for potential pseudogenes, distinct LRR-RKs and wrongly annotated genes using the following filters: A) candidate sequence with a sequence length > 90 % and < 110 % of Arabidopsis *MIK2* were retained, B) sequences with gaps or inserts within strongly conserved domains were removed. Initial alignments were performed using the online version of MAFFT 7 using the E-INS-i strategy (Dataset S3), the L-INS-I strategy was preferred once major gaps were removed in the alignment (68). A phylogenetic analysis was performed on the CIPRES web portal using RAXML-HPC2 on XSEDE (v8.2.12) with the automatic protein model assignment algorithm using ML criterion and 250 bootstrap replicates (69, 70). The JTT likelihood protein model was selected as the best scoring model for ML analysis. The resulting tree was rooted using an LRR-RK found at the conserved MIK2 locus in *T. cacao*, visualized using MEGA10, and edited in Corel-DRAW Home & Student x7. The above strategy was repeated with just the extracellular domains (71).

### RCM

RCM plots were created as previously described (48). In short, we extracted the LRR domains with phyto-predictlrr (obtained in December 2021) (72), aligned them with muscle (version 3.8.31, (62)), trimmed gaps with trimal (version v1.4.rev22) (73), calculated conservation scores with mstatx (1-weighted entropy, github.com/gcollet/MstatX), and extracted a consensus sequence with em_cons (74). To plot the conservation score onto the potential three-dimensional structure, positions of the LRR repeats were again identified with phyto-predictlrr.

#### Brassicales species tree

The tree shown in Fig. S1 was extracted from a larger tree that spanned the 350 plant species previously described(46, 75). To construct the original phylogenetic tree with 350 species, we adapted a previously described protocol (76). First, we searched single-copy genes with BUSCO using the viridiplantae_odb10 database (version 5.5.0) (77). No gene from the BUSCO database was found in all species (the maximum was 300 species at once). We thus used 425 BUSCO-genes that occurred in more than 200 species. For each gene, we extracted all sequences, aligned them with muscle (62), trimmed the alignment with trimal (version 1.4.rev22, option -gt 0.9) (73), and constructed a tree with FastTree (version 2.1.11 SSE3, option - lg) (66). Trees from all genes were finally merged into a single tree that included all species using ASTRAL (version 5.6.3) (78). The final tree was rooted with *C. paradoxa*. Subsequently, the tree was converted to an ultrametric tree using the function chronos() from the R-package “ape” (version 5.6-2) (78). Ultimately, for Fig. S1, we extracted the branch that includes the Brassicales as *PROSCOOPs* are limited to this order and retained *C. papaya* as the closest outgroup.

#### Plant materials and synthetic peptides

*Arabidopsis thaliana* ecotype Columbia (Col-0) was used as wild-type control for SCOOP-induced ROS production and were, similar to other *Brassicales*, grown in growth chambers (20 °C, 60 % RH and 10:14 light:dark cycles). Other plants tested were: *Carica papaya* Tainung, *Cleome violacea* N29053, *Eutrema salsugineum* N22504, *Euclidium syriacum* GC 0587-68, *Diptychocarpus strictus* KM 05-0397-10-00, *Brassica rapa* R500. *Nicotiana benthamiana* was used for experiments which leveraged heterologous expression and were grown in the greenhouse (25/22 °C day/night, 60 % RH and 16:8 light:dark cycles). All synthetic peptides were ordered at >80 % purity (physiological assays) or >95 % purity (biochemical assays) (EZBiolabs) and were earlier described (17).

#### SCOOP-induced ROS production within and outside the Brassicales

All peptides used in this study were synthesized and reconstituted in H_2_O. SCOOP12 (PVRSSQSSQAGGR), SCOOP16 (YVPPSKSRRGKGP), SCOOP21 (YVPPSKSRRGKGP) and SCOOP24 (RVPRSKSPPDRQW) were leveraged to test the activity of putative MIK2 homologues. The flg22 peptide (sequence QRLSTGSRINSAKDDAAGLQIA) originates from bacterial flagellin and was used as a positive control for ROS production upon peptide treatment (79). Leaf punches were taken with a 4-mm biopsy punch and floated in 100 μL of H_2_O using individual cells of a white 96-well white bottom plate (Greiner F-Boden, lumitrac, med. Binding, [REF 655075]). After overnight incubation, H_2_O was removed and ROS production was measured upon addition of a 100-μL assay solution which contained 10 μg/mL horseradish peroxidase (P6782, Merck), 10 mM luminol and the treatment (1 μM SCOOP, 0.1 μM flg22 or H_2_O). Luminescence was quantified with a HIGH-RESOLUTION PHOTON COUNTING SYSTEM (HRPCS218, Photek). Four technical replicates per biological replicate (specified in Fig. S2) were quantified for each treatment, significant differences were determined by performing a two-group Mann-Whitney U Test between each SCOOP treatment and the mock treatment. R and the R-packages dplyr (v1.1.2), ggpubr (v.0.6.0), and ggplot2 (v3.4.2) were used to analyze and plot the data. The resulting figure was edited in Corel-DRAW Home & Student x7.

### Functional validation of MIK2 homologues

#### Molecular cloning

All constructs were created using a hierarchical modular cloning approach facilitated by the MoClo toolkit and the MoClo Plant Parts kit (80, 81). For recombinant expression, we used the previously published Arabidopsis *MIK2* sequence (L0 level backbone, CZLp4057) (18), and *MIK2* homologues were synthesized with domesticated BsaI and BpiI sites and inserted in a pMA-RQ backbone (Invitrogen, Thermo Fisher Scientific). Subsequently, the L0 fragments and an mEGFP C-terminal tag (CZLp4772) were inserted into level 1 Golden-Gate plasmids CZLp4130, which already includes a 35S promotor (CaMV) and an OCS terminator. GoldenGate reactions were performed with 5 U of restriction enzyme, 200 U of T4 ligase in T4 ligase buffer (NEB), 0.1 mg/mL BSA (NEB) and 40 GoldenGate digestion ligation cycles (80). All constructs were validated by Sanger sequencing upon completion and plasmid maps can be found in Dataset S4 (Eurofins genomics).

#### Transient expression in *Nicotiana benthamiana*

*N. benthamiana* does not respond to SCOOP12, allowing the use of heterologous expression in *N. benthamiana* to test putative MIK2 function (14). *Agrobacterium tumefaciens* strain GV3101 transformed with the appropriate construct were grown overnight in LB-media and spun-down. The bacteria were resuspended in infiltration media (10 mM MES-KOH, pH 5.8, 10 mM MgCl_2_) and adjusted to an OD_600_ of 0.5. After 3 h of incubation, the youngest fully expanded leaves of 4-to 5-week-old plants were infiltrated.

#### ROS measurements in *Nicotiana benthamiana*

Following Agrobacterium infiltration for receptor expression (24-48 h), leaf punches were taken with a 4-mm biopsy punch and floated in 100 μL of H_2_O using individual cells of a white 96-well white bottom plate (Greiner F-Boden, lumitrac, med. Binding, [REF 655075]). Subsequently, the same procedure was followed as outlined before while using diverse Arabidopsis SCOOPs (AA sequences, Dataset S1) and flg22. Biological replicates were quantified (n≥4 plants), with each biological replicate representing four technical replicates. R and the R-packages dplyr (v1.1.2), ggpubr (v.0.6.0), and ggplot2 (v3.4.2) were used to analyze and plot the data. The resulting figure was edited in Corel-DRAW Home & Student x7.

#### Cytoplastic calcium measurements in *Nicotiana benthamiana*

Following Agrobacterium infiltration for receptor expression (24 h) in a stable aequorin expressing line of *N. benthamiana* (82), leaf punches were taken with a 4-mm biopsy punch and floated in 100 μL of H_2_O with 20 μM coelenterazine (Merck), using individual cells of a white 96-well white bottom plate (Greiner F-Boden, lumitrac, med. Binding, [REF 655075]). After overnight incubation, the coelenterazine solution was replaced with 100 μL H_2_O and rested for a minimum of 30 min in the dark. Two readings were taken in a TECAN SPARK plate reader every minute for 45 min using an integration time of 250 ms. Biological replicates were quantified (n=4 plants), with each biological replicate representing four technical replicates. R and the R-packages dplyr (v1.1.2), ggpubr (v.0.6.0), and ggplot2 (v3.4.2) were used to analyze and plot the data. The resulting figure was edited in Corel-DRAW Home & Student x7.

#### Protein extraction and western blotting

*N. benthamiana* leaf tissues were flash-frozen in liquid nitrogen and grounded using plastic pestles in 1.5-mL microcentrifuge tubes. Grounded tissue was mixed with 2× loading sample buffer (4 % SDS, 20 % glycerol, 20 mM DTT, 0.004 % bromophenol blue, and 100 mM Tris-HCl pH 7.5) for 10 min at 95 °C. Subsequently, samples were spun at 13,000 × g for 2 min prior to loading and running on a 1.5-mm 10 % SDS-PAGE gels. Proteins were transferred onto PVDF membrane (ThermoFisher) prior to incubation with α-GFP (B-2) HRP (Santa Cruz 9996 HRP, 1:1500). Western blots were imaged with a Bio-Rad ChemiDoc and Image Lab Touch Software (v2.2.0.08). Protein loading was visualized by staining the blotted membrane with Coomassie brilliant blue.

#### AlphaFold-Multimer (AFM)

Protein structure complex predictions of the extracellular domain MIK2 (AT4G08850) with all 50 Arabidopsis SCOOPs were created using the ColabFold platform (v1.3.0) (17, 83). The extracellular domain of MIK2 was determined using deepTMHMM (71). The AF-multimer input sequence alignment was generated through MMseqs2 using the unpaired+paired mode without using templates(83–86). Three recycles were run for each of the five created models. The five resulting models were ranked based by AFM (0.8^*^ipTM + 0.2^*^predicted Template Modelling (pTM) score). The structure files (.pdb) of the twelve highest scoring complexes and their corresponding predicted aligned error (PAE) files are provided in Dataset S5. (87)R and the R-packages dplyr (v1.1.2), ggpubr (v.0.6.0), and ggplot2 (v3.4.2) were used to analyze and plot the ipTM data. The resulting figure was edited in Corel-DRAW Home & Student x7.

#### Mutagenesis

All primers and plasmids used and generated in this study are listed (Table S1, Dataset S4). Site-directed mutagenesis (SDM) was conducted as described by (88). The L0 construct of Arabidopsis *MIK2* was used as template (CZLp4057 (18)). The PCR reaction was DpnI (New England Biolabs) digested at 37 °C for 2 h without prior clean-ups, and then transformed into *E. coli* DH10b. Similar as described before, the L1 constructs were completed by insertion into a level 1 Golden-Gate plasmid CZL4130, which already includes a 35S promotor (CaMV) and a NOS terminator, and the addition of an mEGFP C-terminal tag (CZLp4772). GoldenGate reactions were performed with 5 U of restriction enzyme, 200 U of T4 ligase in T4 ligase buffer (NEB), 0.1 mg/mL BSA (NEB) and 40 GoldenGate digestion ligation cycles (80). All constructs were validated by whole plasmid sequencing, plasmid maps can be found in Dataset S4 (Eurofins genomics).

#### Protein expression and purification

*Spodoptera frugiperda* codon-optimized synthetic genes (Invitrogen GeneArt), coding for *Arabidopsis thaliana* MIK2 ectodomain (residues 1 to 709) mutants were cloned into a modified pFastBAC vector (Geneva Biotech) with its native signal peptide, a C-terminal TEV (tobacco etch virus protease) cleavable site and a StrepII-9xHis affinity tag. Baculovirus generation was carried out using DH10 cells and virus production and amplification was done in Sf9 cells. *Trichoplusia ni* Tnao38 cells were used for protein expression (89), that were infected with MIK2 mutant viruses with a multiplicity of infection (MOI) of 3 and incubated 1 day at 28°C and 2 days at 22°C at 110 rpm. The secreted proteins and complexes were purified by Ni^2+^ (HisTrap excel, Cytiva, equilibrated in 25 mM KP_i_ pH 7.8 and 500 mM NaCl) followed by Strep (Strep-Tactin Superflow high-capacity, IBA Lifesciences, equilibrated in 25 mM Tris pH 8.0, 250 mM NaCl, 1 mM EDTA) affinity chromatography. All proteins were incubated with TEV protease to remove the tags. Proteins were further purified by SEC on a Superdex 200 Increase 10/300 GL column (Cytiva) equilibrated in 20 mM citric acid pH5.0, 150 mM NaCl. Proteins were concentrated using Amicon Ultra concentrators (Millipore, molecular weight cut-off 3,000, 10,000 and 30,000), and SDS-PAGE was used to assess the purity and structural integrity of the different proteins.

#### Isothermal titration calorimetry (ITC)

Experiments were performed at 25 °C using a MicroCal PEAQITC (Malvern Instruments) with a 200 μL standard cell and a 40 μL titration syringe. The MIK2 mutant ectodomains were gel filtrated into pH 5 ITC buffer (20 mM citric acid pH 5.0, 150 mM NaCl). SCOOP12 peptide powder was dissolved in the same buffer to obtain the desired concentration. A typical experiment consisted of injecting 3 μL of a 150 or 300 μM solution of the ligand into 15 μM MIK2 solution in the cell at 150 s intervals and 500 rpm stirring speed. ITC data were corrected for the heat of dilution by subtracting the mixing enthalpies for titrant solution injections into protein-free ITC buffer. Experiments were done in duplicates and data were analyzed using the MicroCal PEAQ-ITC Analysis Software provided by the manufacturer. All ITC runs used for data analysis had an N ranging between 0.8 and 1.3. The N values were fitted to 1 in the analysis.

#### Analytical size-exclusion (SEC) chromatography

Analytical SEC experiments were performed using a Superdex 200 Increase 10/300 GL column (GE). The columns were pre-equilibrated in 20 mM citric acid pH 5, 150 mM NaCl. One hundred fifty micrograms of MIK2 mutant ectodomains were injected sequentially onto the column and eluted at 0.5 mL/min. Ultraviolet absorbance (UV) at 280 nm was used to monitor the elution of the proteins. The peak fractions were analyzed by SDS-PAGE followed by Coomassie blue staining.

#### Structural visualization and model analysis

MIK2-SCOOP prediction models obtained from AFM were superimposed using UCSF Chimera (90). Molecular diagrams have prepared with PyMOL (87), retrieved from http://www.pymol.org/pymol).

#### Co-immunoprecipitation (Co-IP)

Following Agrobacterium infiltration of receptor (variants), BAK1 (CZLp3593) and P19 (CZLp5085) expression (48 h), leaves were split in half and midveins were removed. This way, mock- and treatment of interest can later be performed on two samples created from the same infiltration event. All samples were submerged in 0.25x MS-sucrose for 30 min. Next, samples were vacuum infiltrated with 1 μM SCOOP12 in 0.25x MS-sucrose or just 0.25x MS-sucrose as a mock treatment. Finally, all samples were dried with paper towel and flash frozen.

For co-immunoprecipitation assays, approximately 3.5 g of frozen tissue was ground to a fine powder in nitrogen-cooled stainless-steel jars using a Retsch MM300 ball mill. Tissue was thawed in extraction buffer (50 mM Tris-HCl pH 7.5, 150 mM NaCl, 2 mM EDTA, 10 % (v/v) glycerol, 2 mM DTT and 1:100 home-made protease inhibitor cocktail equivalent to Sigma-Aldrich P9599) at a ratio of 2 ml of buffer per gram of tissue and proteins were solubilized on a rotator at 4°C for 30 min. Extracts were filtered through two layers of Miracloth and centrifuged at 25,000 x g for 30 min at 4 °C to generate a clarified extract. Protein amounts were estimated using the Bradford assay and samples were normalized to contain equal amounts of protein.

Protein extracts containing GFP-tagged MIK2 or site-directed mutants were incubated with 20 µL of GFP-Trap beads (Chromotek) for 2 h with gentle mixing at 4 °C to immuno-precipitate receptor complexes. The beads were sedimented by centrifugation at 1000 x g for 4 min at 4 °C and were subsequently suspended in 1 mL of extraction buffer (see above). The beads were sedimented at 1,000 x g for 1 min and suspended in 1 mL of extraction buffer three more times for a total of four washes. After the last wash was removed, beads were suspended in 2X Laemmli SDS-PAGE loading buffer followed by heating at 80 °C for 10 min. Five microliters of each IP fraction was loaded into an 8 % (v/v) SDS-PAGE gel and proteins were separated for 90 min at 150 V followed by transfer to PVDF membrane for immunoblotting with anti-GFP (B-2) and anti-BAK1 antibodies.

## Supporting information

Snoeck_SI

## Acknowledgements

This research was supported by the University of Zurich and the European Research Council under Grant Agreement No. 773153 (‘IMMUNO-PEPTALK’ to CZ), and by the University of Lausanne and the Swiss National Science Foundation (grant 310030_204526 to JS). ADFF is also supported by a Post-Doctoral Fellowship from the European Molecular Biology Organization (EMBO, ALTF 580-2022). We thank Owen Kentish for assistance with recombinant protein production. We thank members of the Zipfel lab for discussion during this project and feedback on the manuscript.

## Supporting information legends

Fig. S1: **A species phylogeny that indicates besides the presence of putative (PRO)SCOOP- and MIK2-homologues also SCOOP-induced plant-signaling responses in native plants and post heterologous expression (HE) in *N. benthamiana***.

Fig. S2: **Diverse species of the order of the Brassicales respond to SCOOP treatment with reactive oxygen species (ROS) production. A)** Shown is ROS production in cumulative relative luminescence units (RLU) for (4-30 min or 4-60 min), four technical replicates per biological replicate (# indicated in the figure), in relative luminescence units (RLUs) (1 observation/min) after treatment with H_2_O (white), SCOOP12, SCOOP13, SCOOP16 and SCOOP24 (1 μM, grey) or the peptide flg22 (1 μM, dark grey). Significant differences between the control and the treatments of interest were found by performing a Wilcoxon rank-sum test. **B)** Sequence motif analysis of SCOOP13-16 and SCOOP24. Sequence logos were generated using Dataset S1 and WebLogo server (https://weblogo.berkeley.edu/logo.cgi).

Fig. S3: **SCOOP-dependent reactive oxygen species (ROS) production and Ca**^**2+**^ **influx following the heterologous expression of MIK2 and MIK2 homologues in *N. benthamiana*. A-B)** Shown are ROS production (A) and Ca^2+^ influx (B), respectively 4-60 min and 3-45 min, in cumulative relative luminescence units (RLUs) post treatment with H_2_O (white) or SCOOP12, SCOOP13, SCOOP16 and SCOOP24 (1 μM, grey). Each biological replicate (n=4 plants) is represented by four technical replicates. Significance was tested by performing a paired Wilcoxon rank-sum test.

Fig. S4: **Confocal microscopy and western following the heterologous expression of MIK2 and MIK2 variants in *N. benthamiana*. A)** Western blot 72 h post-Agrobacterium infiltration. The western blot was probed with α-GFP (B-2) HRP as the receptor had a C-terminal GFP tag (top) and subsequently stained with CBB as a loading control (bottom). **B)** Confocal microscopy (GFP, Chlorophyll B and Bright Field) following Agrobacterium infiltration (72 h). All confocal microscopy images were identically modified, with small adjustments of brightness and contrast. The scale bar represents 20 μm. Plasmolysis was obtained by treatment with 0.6 M mannitol for 30 min. A repeat was performed and confirmed the depicted results.

Fig. S5. **Analytical size-exclusion chromatography experiments (SEC) of MIK2 pocket variants. SDS-PAGE of the proteins eluted are presented alongside**.

Fig. S6. **ITC assays of MIK2 pocket variants and SCOOP12. ITC thermograms of the independent experiments performed for each mutant and analyzed in Fig. 4C**.

Fig. S7: **SCOOP-dependent reactive oxygen species (ROS) production following the heterologous expression of MIK2 and MIK2 variants in *N. benthamiana***. At least four independent biological replicates were performed (n≥4 plants), with each biological replicate represented by four technical replicates. Cumulative RLUs representing ROS production are shown after treatment with H_2_O (white) or the SCOOP peptide indicated (1 μM, grey). Significance was tested by performing a paired Wilcoxon rank-sum test.

Table S1: **Primers used in this study**.

Dataset S1: **Overview SCOOP mining**.

Dataset S2: **Overview of mined assemblies for locus analyses, overview of Arabidopsis anchor genes *PROSCOOP* loci, and overview of contiguous *INR* loci (+ coordinates) and MIK2-orthologues**.

Dataset S3: **Fasta file of the amino acid (AA) sequences of MIK2 homologues, putative MIK2 paralogues and outgroup included in the phylogenetic analysis**.

Dataset S4: **Plasmid maps of constructs used in this study**.

Dataset S5: **AFM predicted structures (.pdb) and confidence metrics (.pae)**

Dataset S6: **Unedited files (.tiff) of co-IP and western blotting**.

## References

1. V. Olsson, et al., Look closely, the beautiful may be small: precursor-derived peptides in plants. Annu. Rev. Plant Biol 70, 153–186 (2019).

2. C. Furumizu, et al., The sequenced genomes of nonflowering land plants reveal the innovative evolutionary history of peptide signaling. Plant Cell 33, 2915–2934 (2021).

3. H. Zhang, X. Lin, Z. Han, L.-J. Qu, J. Chai, Crystal structure of PXY-TDIF complex reveals a conserved recognition mechanism among CLE peptide-receptor pairs. Cell Res 26, 543–555 (2016).

4. J. Tang, et al., Structural basis for recognition of an endogenous peptide by the plant receptor kinase PEPR1. Cell Res 25, 110–120 (2015).

5. J. Santiago, et al., Mechanistic insight into a peptide hormone signaling complex mediating floral organ abscission. Elife 5 (2016).

6. Y. Xiao, et al., Mechanisms of RALF peptide perception by a heterotypic receptor complex. Nature 572, 270–274 (2019).

7. A. O. Roman, et al., HSL1 and BAM1/2 impact epidermal cell development by sensing distinct signaling peptides. Nature Communications 2022 13:1 13, 1–13 (2022).

8. W. Song, et al., Signature motif-guided identification of receptors for peptide hormones essential for root meristem growth. Cell Res 26, 674–685 (2016).

9. X. Zhang, et al., Structural basis for receptor recognition of pollen tube attraction peptides. Nat Commun 8, 1–9 (2017).

10. S. Okuda, et al., Molecular mechanism for the recognition of sequence divergent CIF peptides by the plant receptor kinases GSO1/SGN3 and GSO2. Proc Natl Acad Sci U S A 117, 2693–2703 (2020).

11. J. Wang, et al., Allosteric receptor activation by the plant peptide hormone phytosulfokine. Nature 525, 265–268 (2015).

12. P. Bryant, et al., Predicting the structure of large protein complexes using AlphaFold and Monte Carlo tree search. Nat Commun 13, 1–14 (2022).

13. M. A. Outram, M. Figueroa, J. Sperschneider, S. J. Williams, P. N. Dodds, Seeing is believing: exploiting advances in structural biology to understand and engineer plant immunity. Curr Opin Plant Biol 67, 102210 (2022).

14. K. Gully, et al., The SCOOP12 peptide regulates defense response and root elongation in Arabidopsis thaliana. J Exp Bot 70, 1349–1365 (2019).

15. S. Hou, et al., The Arabidopsis MIK2 receptor elicits immunity by sensing a conserved signature from phytocytokines and microbes. Nat Commun 12, 1–15 (2021).

16. J. Zhang, et al., EWR1 as a SCOOP peptide activates MIK2-dependent immunity in Arabidopsis. J Plant Interact 17, 562–568 (2022).

17. H. Yang, et al., Subtilase-mediated biogenesis of the expanded family of SERINE RICH ENDOGENOUS PEPTIDES. Nat Plants 9, 2085–2094 (2023).

18. J. Rhodes, et al., Perception of a divergent family of phytocytokines by the Arabidopsis receptor kinase MIK2. Nat Commun 12, 1–10 (2021).

19. A. Dievart, C. Gottin, C. Périn, V. Ranwez, N. Chantret, Origin and diversity of plant receptor-like kinases. Annu Rev Plant Biol 71, 131–156 (2020).

20. U. Hohmann, K. Lau, M. Hothorn, The structural basis of ligand perception and signal activation by receptor kinases. Annu Rev Plant Biol 68, 109–137 (2017).

21. D. Restrepo-Montoya, R. Brueggeman, P. E. McClean, J. M. Osorno, Computational identification of receptor-like kinases “RLK” and receptor-like proteins “RLP” in legumes. BMC Genomics 21, 1–17 (2020).

22. I. Albert, C. Hua, T. Nürnberger, R. N. Pruitt, L. Zhang, Surface sensor systems in plant immunity.Plant Physiol 182, 1582–1596 (2020).

23. I. Fischer, A. Diévart, G. Droc, J.-F. Dufayard, N. Chantret, Evolutionary dynamics of the leucine-rich repeat receptor-like kinase (LRR-RLK) subfamily in angiosperms. Plant Physiol 170, 1595–1610 (2016).

24. D. Van der Does, et al., The Arabidopsis leucine-rich repeat receptor kinase MIK2/LRR-KISS connects cell wall integrity sensing, root growth and response to abiotic and biotic stresses. PLoS Genet 13 (2017).

25. M. C. Guillou, et al., The peptide SCOOP12 acts on reactive oxygen species homeostasis to modulate cell division and elongation in Arabidopsis primary root. J Exp Bot 73, 6115–6132 (2022).

26. E. Stahl, et al., The MIK2/SCOOP signaling system contributes to Arabidopsis resistance against herbivory by modulating jasmonate and indole glucosinolate biosynthesis. Front Plant Sci 13, 852808 (2022).

27. M. C. Guillou, et al., The PROSCOOP10 gene encodes two extracellular hydroxylated peptides and impacts flowering time in Arabidopsis. Plants 11, 3554 (2022).

28. Z. Zhang, N. Gigli-Bisceglia, W. Li, C. Testerink, Y. Guo, Antagonistic regulation of Arabidopsis leaf senescence by SCOOP10 and SCOOP12 peptides via MIK2 receptor-like kinase. bioRxiv, 2023.10.27.564453 (2023).

29. A. D. Coleman, et al., The Arabidopsis leucine-rich repeat receptor-like kinase MIK2 is a crucial component of early immune responses to a fungal-derived elicitor. New Phytologist 229, 3453–3466 (2021).

30. J. Zhang, et al., EWR1 as a SCOOP peptide activates MIK2-dependent immunity in Arabidopsis. J Plant Interact 17, 562–568 (2022).

31. S. Moussu, J. Santiago, Structural biology of cell surface receptor-ligand interactions. Curr Opin Plant Biol 52, 38–45 (2019).

32. H. Zhang, Z. Han, W. Song, J. Chai, Structural insight into recognition of plant peptide hormones by receptors. Mol. Plant 9, 1454–1463 (2016).

33. S. Snoeck, et al., Evolutionary gain and loss of a plant pattern-recognition receptor for HAMP recognition. Elife 11, e81050 (2022).

34. L. Zhang, et al., Distinct immune sensor systems for fungal endopolygalacturonases in closely related Brassicaceae. Nat Plants 7, 1254–1263 (2021).

35. Y. C. Torres Ascurra, et al., Functional diversification of a wild potato immune receptor at its center of origin. Science 381, 891–897 (2023).

36. U. Fürst, et al., Perception of Agrobacterium tumefaciens flagellin by FLS2XL confers resistance to crown gall disease. Nature Plants 2020 6:1 6, 22–27 (2020).

37. Y. Wei, et al., An immune receptor complex evolved in soybean to perceive a polymorphic bacterial flagellin. Nature Communications 2020 11:1 11, 1–11 (2020).

38. R. Evans, et al., Protein complex prediction with AlphaFold-Multimer. BioRxiv, 1–25 (2022).

39. P. Bryant, G. Pozzati, A. Elofsson, Improved prediction of protein-protein interactions using AlphaFold2. Nat Commun 13, 1–11 (2022).

40. S. Shanker, M. F. Sanner, Predicting protein™peptide interactions: benchmarking deep learning techniques and a comparison with focused docking. J. Chem. Inf. Model 63, 3170 (2023).

41. I. Johansson-Åkhe, B. Wallner, Improving peptide-protein docking with AlphaFold-Multimer using forced sampling. Frontiers in bioinformatics 2, 1–17 (2022).

42. N. B. Danneskiold-Samsøe, et al., AlphaFold2 enables accurate deorphanization of ligands to single-pass receptors. bioRxiv, 1–42 (2023).

43. M. F. Lensink, et al., Impact of AlphaFold on structure prediction of protein complexes: The CASP15-CAPRI experiment. Proteins: Structure, Function and Bioinformatics 91, 1658–1683 (2023).

44. T. Tsaban, et al., Harnessing protein folding neural networks for peptide-protein docking. Nat Commun 13, 1–12 (2022).

45. J. Ko Lee, Can AlphaFold2 predict protein-peptide complex structures accurately? BioRxiv, 1–6 (2021).

46. B. P. M. Ngou, R. Heal, M. Wyler, M. W. Schmid, J. D. G. Jones, Concerted expansion and contraction of immune receptor gene repertoires in plant genomes. Nat Plants 8, 1146–1152 (2022).

47. K. P. Hendriks, et al., Global Brassicaceae phylogeny based on filtering of 1,000-gene dataset. Current Biology 33, 4052–4068 (2023).

48. L. Helft, et al., LRR Conservation mapping to predict functional sites within protein leucine-rich repeat domains. PLoS One 6, e21614 (2011).

49. S. H. Shiu, A. B. Bleecker, Receptor-like kinases from Arabidopsis form a monophyletic gene family related to animal receptor kinases. Proceedings of the National Academy of Sciences 98, 10763–10768 (2001).

50. C. Furumizu, S. Sawa, The RGF/GLV/CLEL family of short peptides evolved through lineage-specific losses and diversification and yet conserves its signaling role between vascular plants and Bryophytes. Front Plant Sci 12, 703012 (2021).

51. J. Rhodes, et al., Perception of a conserved family of plant signalling peptides by the receptor kinase HSL3. Elife 11, e74687 (2022).

52. M. Lori, et al., Evolutionary divergence of the plant elicitor peptides (Peps) and their receptors: interfamily incompatibility of perception but compatibility of downstream signalling. J Exp Bot 66, 5315 (2015).

53. X.-S. Yu, et al., Structure and functional divergence of PIP peptide family revealed by functional studies on PIP1 and PIP2 in Arabidopsis thaliana. Front. Plant Sci 14, 1208549 (2023).

54. Z. Liu, et al., Phytocytokine signalling reopens stomata in plant immunity and water loss. 332 | Nature | 605 (2022).

55. L. Wang, et al., The systemin receptor SYR1 enhances resistance of tomato against herbivorous insects. Nat Plants 4, 152–156 (2018).

56. C. Furumizu, R. B. Aalen, Peptide signaling through leucine-rich repeat receptor kinases: insight into land plant evolution. New Phytologist 238, 977–982 (2023).

57. S. Snoeck, N. Guayazán-Palacios, A. D. Steinbrenner, Molecular tug-of-war: plant immune recognition of herbivory. Plant Cell 34, 1497–1513 (2022).

58. Y. Suzuki, Three-Dimensional window analysis for detecting positive selection at structural regions of proteins. Mol Biol Evol 21, 2352–2359 (2004).

59. C. A. Innis, A. P. Anand, R. Sowdhamini, Prediction of functional sites in proteins using conserved functional group analysis. J Mol Biol 337, 1053–1068 (2004).

60. Y. Ma, et al., Comparisons of two receptor pathways in a single cell-type reveal features of signalling specificity. bioRxiv, 2023.07.03.547518 (2023).

61. M. Steinegger, J. Söding, MMseqs2 enables sensitive protein sequence searching for the analysis of massive data sets. Nat Biotechnol 35, 1026–1028 (2017).

62. R. C. Edgar, MUSCLE: multiple sequence alignment with high accuracy and high throughput. Nucleic Acids Res 32, 1792–1797 (2004).

63. S. R. Eddy, Accelerated profile HMM searches. PLoS Comput Biol 7, 1002195 (2011).

64. A. R. Quinlan, I. M. Hall, BEDTools: a flexible suite of utilities for comparing genomic features. BIOINFORMATICS APPLICATIONS NOTE 26, 841–842 (2010).

65. S. Deorowicz, A. Debudaj-Grabysz, A. Gudyś, FAMSA: Fast and accurate multiple sequence alignment of huge protein families. Sci Rep 6, 1–13 (2016).

66. M. N. Price, P. S. Dehal, A. P. Arkin, FastTree 2 – Approximately maximum-likelihood trees for large alignments. PLoS One 5, e9490 (2010).

67. G. E. Tan, et al., Current methods for automated filtering of multiple sequence alignments frequently worsen single-gene phylogenetic inference. Syst. Biol 64, 778–791 (2015).

68. K. Katoh, K. Misawa, K. Kuma, T. Miyata, MAFFT: a novel method for rapid multiple sequence alignment based on fast Fourier transform. Nucleic Acids Res 30, 3059–3066 (2002).

69. M. A. Miller, W. Pfeiffer, T. Schwartz, Creating the CIPRES science gateway for inference of large phylogenetic trees in 2010 Gateway Computing Environments Workshop, GCE 2010, (IEEE, 2010), pp. 1–8.

70. A. Stamatakis, RAxML version 8: a tool for phylogenetic analysis and post-analysis of large phylogenies. Bioinformatics 30, 1312–3 (2014).

71. J. Hallgren, et al., DeepTMHMM predicts alpha and beta transmembrane proteins using deep neural networks. BioRxiv, 1–12 (2022).

72. T. Chen, Identification and characterization of the LRR repeats in plant LRR-RLKs. BMC Mol Cell Biol 22, 1–16 (2021).

73. S. Capella-Gutiérrez, J. M. Silla-Martínez, T. Gabaldón, trimAl: A tool for automated alignment trimming in large-scale phylogenetic analyses. Bioinformatics 25, 1972–1973 (2009).

74. P. Rice, L. Longden, A. Bleasby, EMBOSS: the European molecular biology open software suite.Trends Genet 16, 276–277 (2000).

75. B. Pok, et al., Evolutionary Trajectory of Pattern Recognition Receptors in Plants. bioRxiv, 2023.07.04.547604 (2023).

76. M. Manni, M. R. Berkeley, M. Seppey, E. M. Zdobnov, BUSCO: assessing genomic data quality and beyond. Curr Protoc 1 (2021).

77. M. Manni, M. R. Berkeley, M. Seppey, F. A. Simão, E. M. Zdobnov, BUSCO update: novel and streamlined workflows along with broader and deeper phylogenetic coverage for scoring of eukaryotic, prokaryotic, and viral genomes. Mol Biol Evol 38, 4647–4654 (2021).

78. C. Zhang, M. Rabiee, E. Sayyari, S. Mirarab, ASTRAL-III: Polynomial time species tree reconstruction from partially resolved gene trees. BMC Bioinformatics 19, 15–30 (2018).

79. G. Felix, J. D. Duran, S. Volko, T. Boller, Plants have a sensitive perception system for the most conserved domain of bacterial flagellin. The Plant Journal 18, 265–276 (1999).

80. E. Weber, C. Engler, R. Gruetzner, S. Werner, S. Marillonnet, A modular cloning system for standardized assembly of multigene constructs. PLoS One 6, e16765 (2011).

81. C. Engler, et al., A Golden Gate modular cloning toolbox for plants. ACS Synth Biol 3, 839–843 (2014).

82. C. Segonzac, et al., Hierarchy and roles of pathogen-associated molecular pattern-induced responses in Nicotiana benthamiana. Plant Physiology Ò 156, 687–699 (2011).

83. M. Mirdita, M. Steinegger, J. Söding, MMseqs2 desktop and local web server app for fast, interactive sequence searches. Bioinformatics 35, 2856–2858 (2019).

84. J. Jumper, et al., Highly accurate protein structure prediction with AlphaFold. Nature 596, 583 (2021).

85. M. Mirdita, et al., Uniclust databases of clustered and deeply annotated protein sequences and alignments. Nucleic Acids Res 45 (2017).

86. A. L. Mitchell, et al., MGnify: the microbiome analysis resource in 2020. Nucleic Acids Res 48, D570–D578 (2020).

87. PyMOL, The PyMOL molecular graphics system, version 2.5.2 Schrödinger, LLC.

88. H. Liu, J. H. Naismith, An efficient one-step site-directed deletion, insertion, single and multiple-site plasmid mutagenesis protocol. BMC Biotechnol 8, 1–10 (2008).

89. Y. Hashimoto, S. Zhang, S. Zhang, Y.-R. Chen, G. W. Blissard, Correction: BTI-Tnao38, a new cell line derived from Trichoplusia ni, is permissive for AcMNPV infection and produces high levels of recombinant proteins. BMC Biotechnol 12, 12 (2012).

90. E. F. Pettersen, et al., UCSF Chimera—A visualization system for exploratory research and analysis.J Comput Chem 25, 1605–1612 (2004).

