## Supplementary material for "Leveraging co-evolutionary insights and AI-based structural modeling to unravel receptor-peptide ligand-binding mechanisms": Snoeck_SI: Snoeck_SI_180124_final for bioRxiv.docx

**This PDF file includes:**

Figures S1 to S7

Table S1

Legends for Datasets S1 to S2

**Other supporting materials for this manuscript include the following:**

Datasets S1 to S2

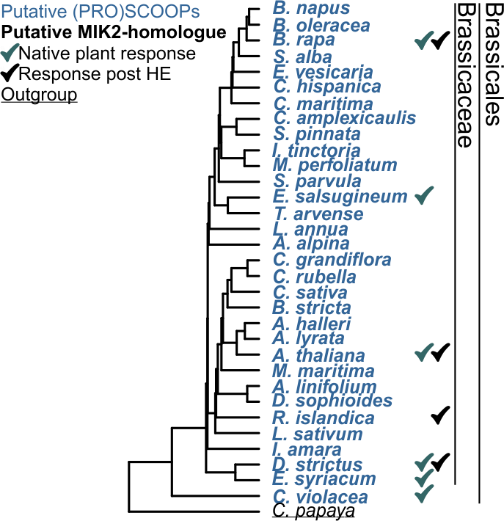

Fig. S1: A species phylogeny that indicates besides the presence of putative SCOOP- and MIK2-homologues also SCOOP-induced plant-signaling responses in native plants and post heterologous expression (HE) in *N. benthamiana.*

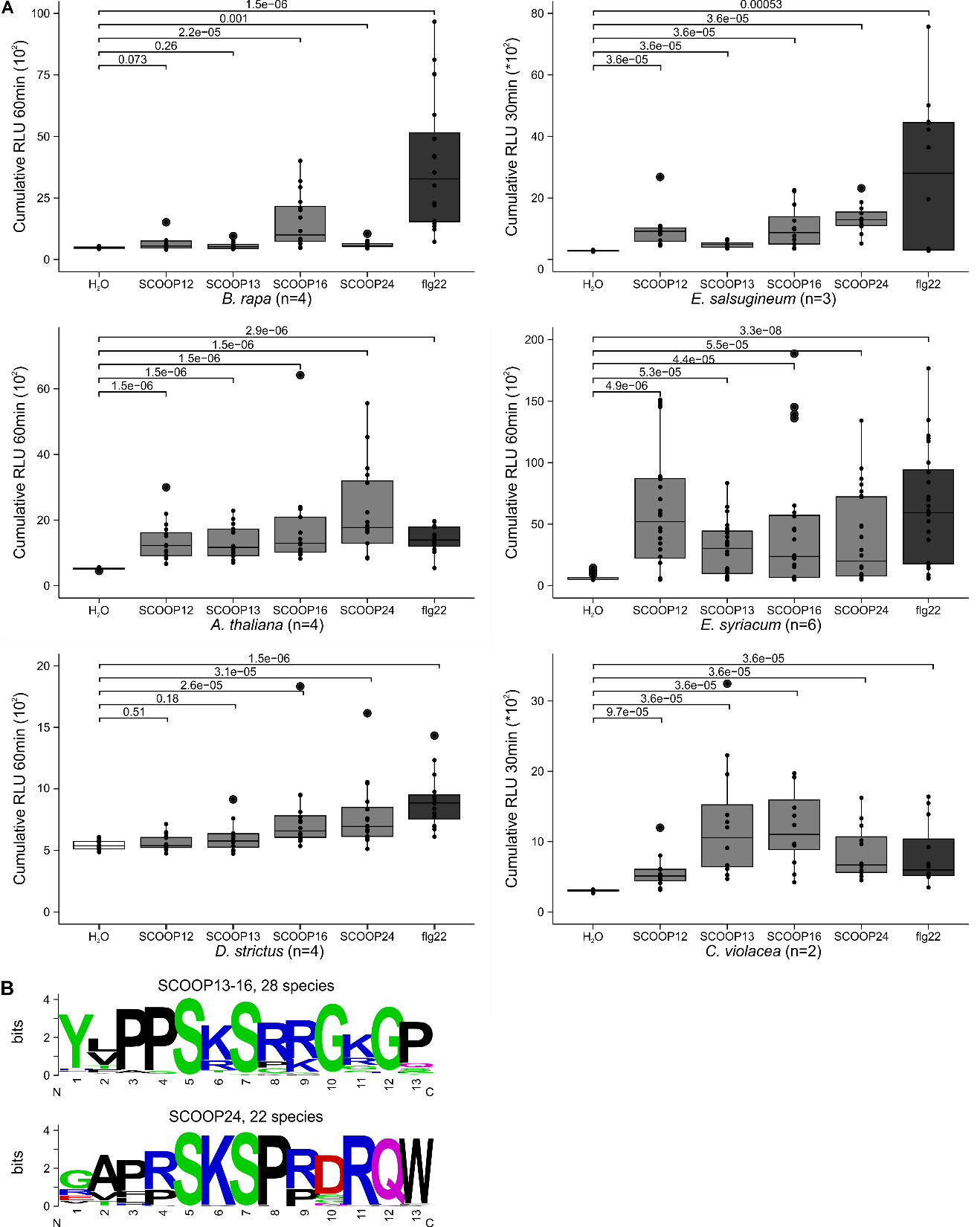

Fig. S2: **Diverse species of the order of the Brassicales respond to SCOOP treatment with reactive oxygen species (ROS) production. A)** Shown is ROS production in cumulative relative luminescence units (RLU) for (4-30 min or 4-60 min), four technical replicates per biological replicate (# indicated in the figure), in relative luminescence units (RLUs) (1 observation/min) after treatment with H_2_O (white), SCOOP12, SCOOP13, SCOOP16 and SCOOP24 (1 μM, grey) or the peptide flg22 (1 μM, dark grey). Significant differences between the control and the treatments of interest were found by performing a Wilcoxon rank-sum test. **B)** Sequence motif analysis of SCOOP13-16 and SCOOP24. Sequence logos were generated using Dataset S1 and WebLogo server (https://weblogo.berkeley.edu/logo.cgi).

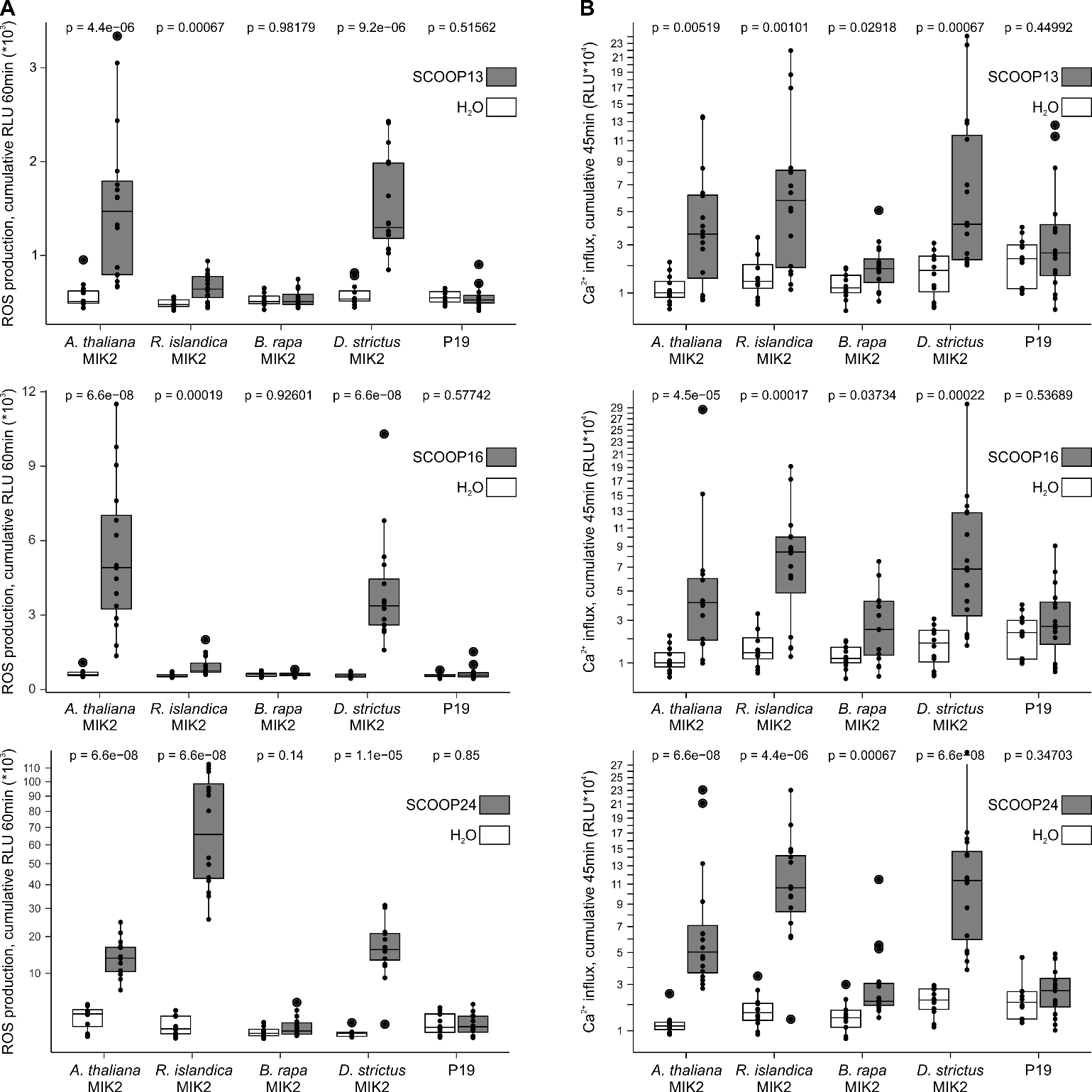

Fig. S3: **SCOOP-dependent reactive oxygen species (ROS) production and Ca^2+^ influx following the heterologous expression of MIK2 and MIK2 homologues in *N. benthamiana*. A-B)** Shown are ROS production (A) and cumulative influx of Ca^2+^ (B), respectively 4-60 min and 3-45 min, in cumulative relative luminescence units (RLUs) post treatment with H_2_O (white) or SCOOP12, SCOOP13, SCOOP16 and SCOOP24 (1 μM, grey). Each biological replicate (n=4 plants) is represented by four technical replicates. Significance was tested by performing a paired Wilcoxon rank-sum test.

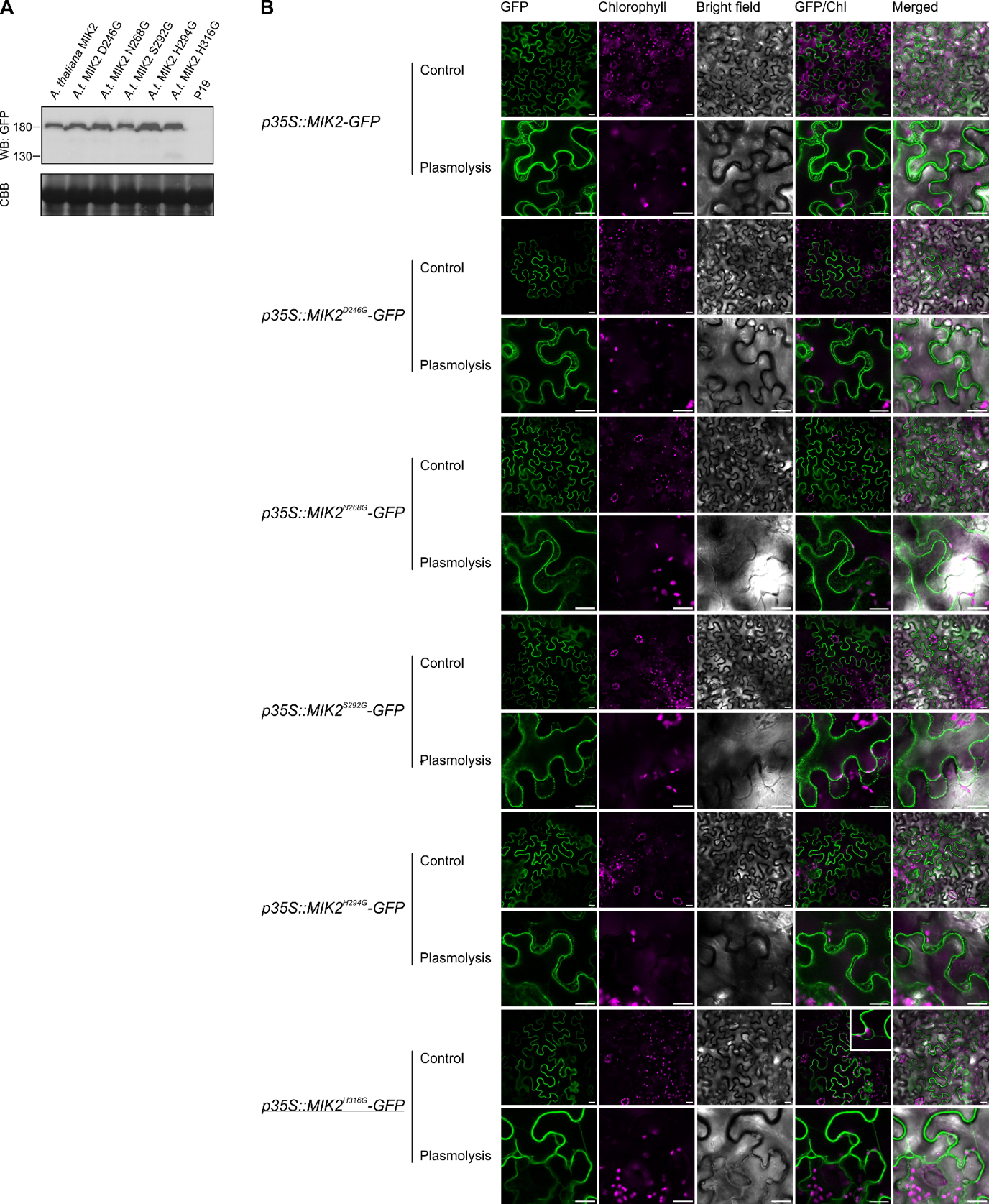

Fig. S4: **Confocal microscopy and western following the heterologous expression of MIK2 and MIK2 variants in *N. benthamiana*. A)** Western blot 72 h post-Agrobacterium infiltration. The western blot was probed with α-GFP (B-2) HRP as the receptor had a C-terminal GFP tag (top) and subsequently stained with CBB as a loading control (bottom). **B)** Confocal microscopy (GFP, Chlorophyll B and Bright Field) following Agrobacterium infiltration (72 h). All confocal microscopy images were identically modified, with small adjustments of brightness and contrast. The scale bar represents 20 μm. Plasmolysis was obtained by treatment with 0.6 M mannitol for 30 min. Raw files can be found in dataset S5. A repeat was performed and confirmed the depicted results.

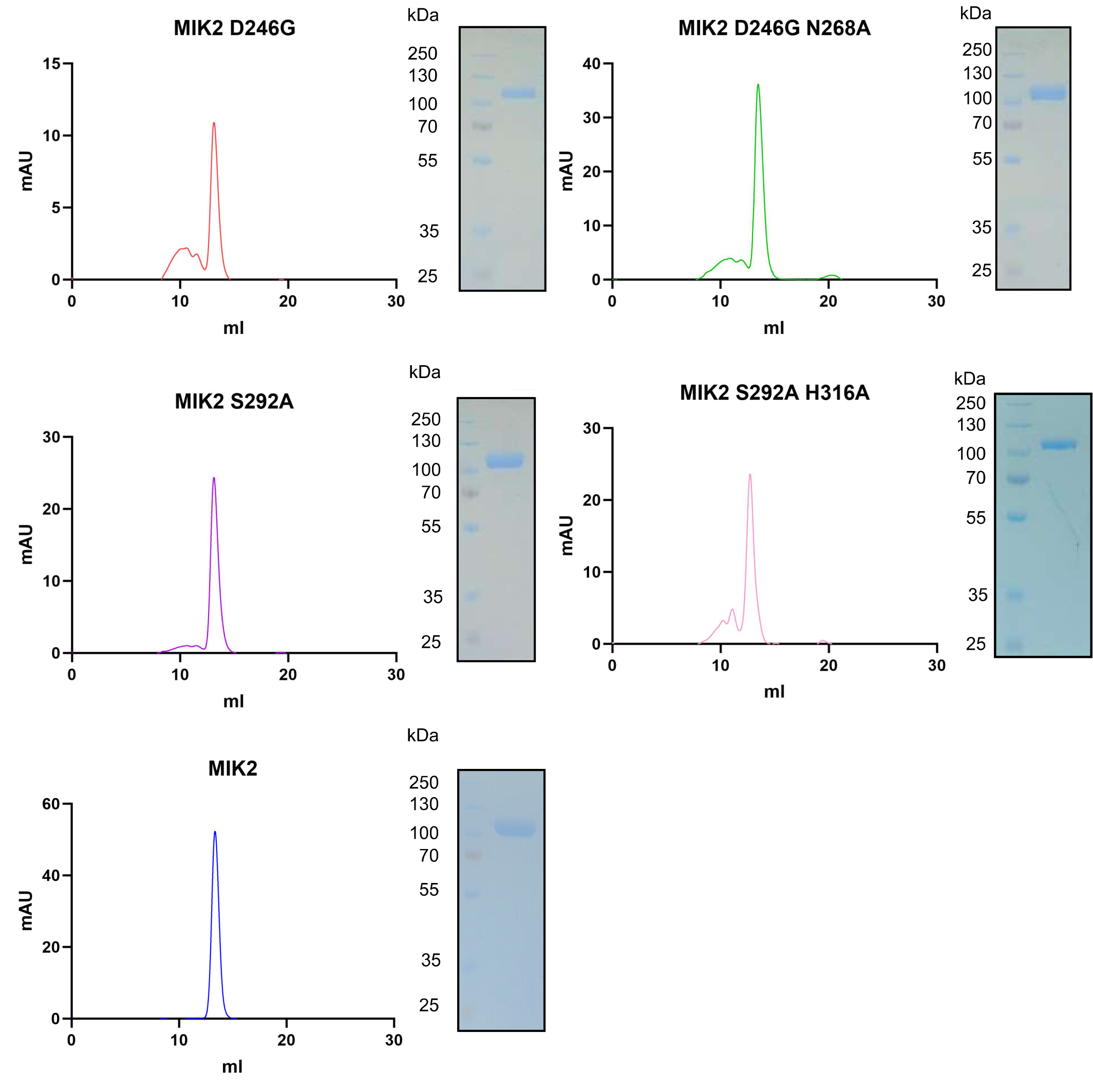
Fig S5. **Analytical size-exclusion chromatography experiments (SEC) of MIK2 pocket variants. SDS-PAGE of the proteins eluted are presented alongside.**

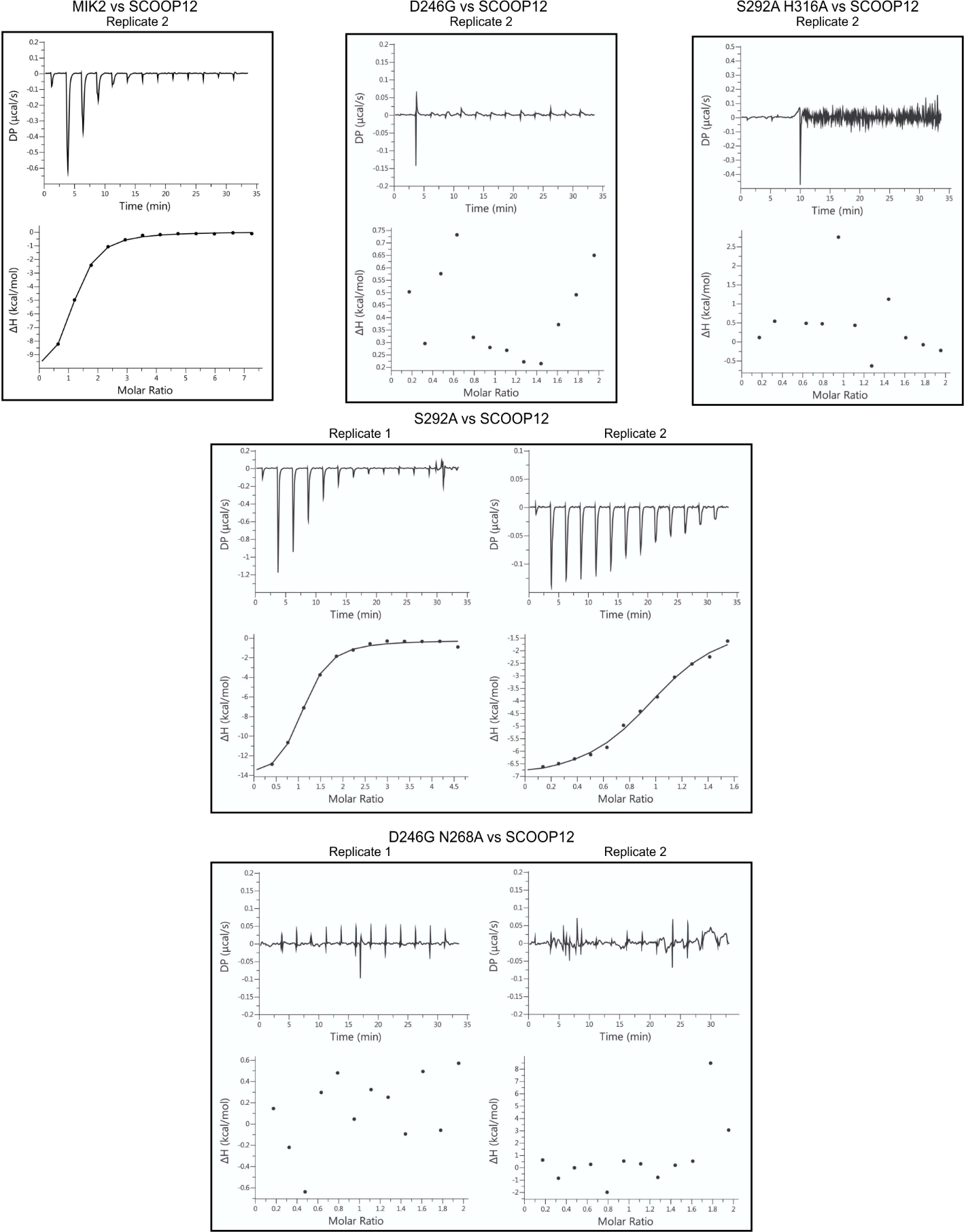

Fig S6. **ITC assays of MIK2 pocket variants and SCOOP12. ITC thermograms of the independent experiments performed for each mutant and analyzed in Figure 4C.**

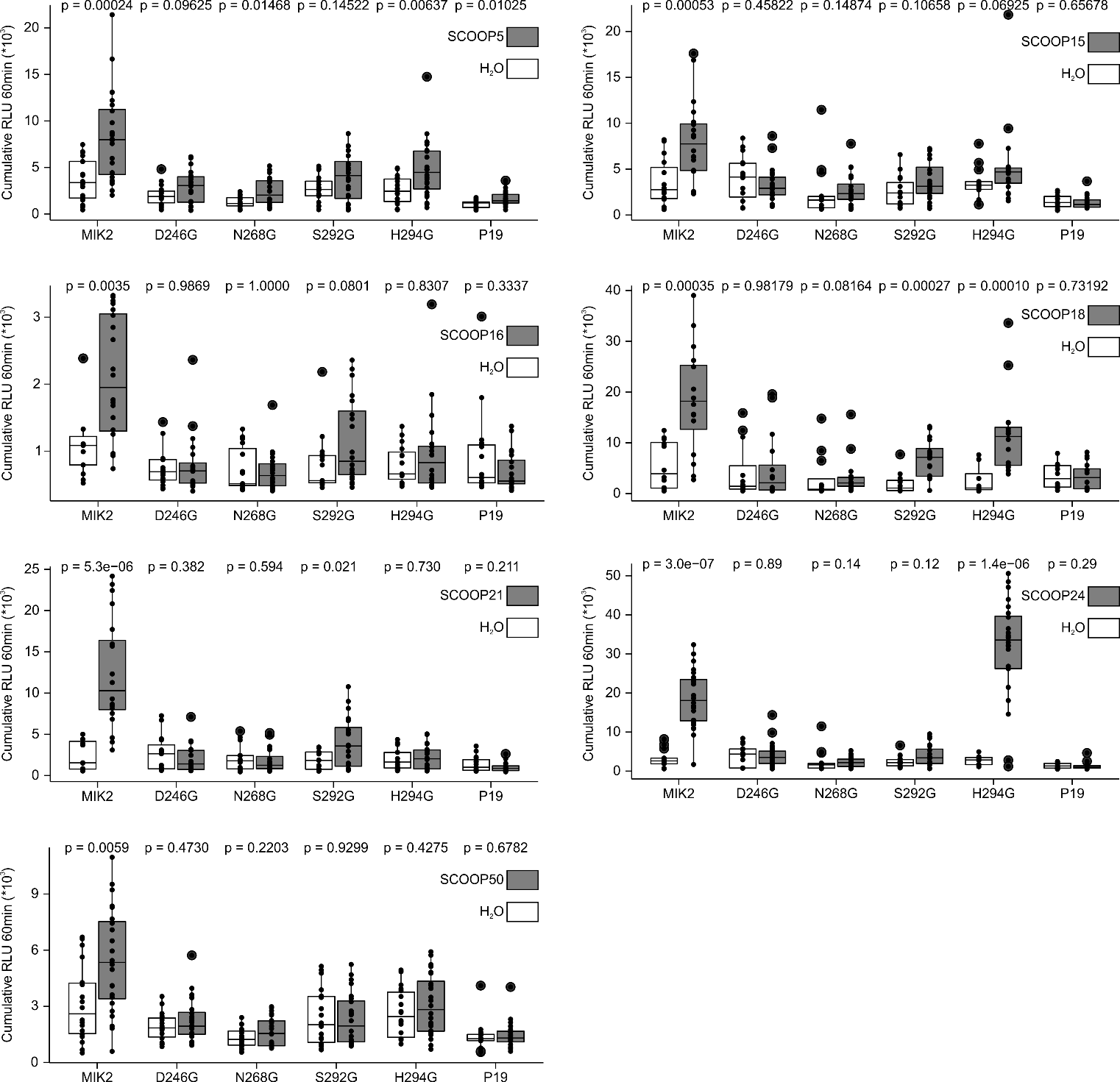

Fig. S7: **SCOOP-dependent reactive oxygen species (ROS) production following the heterologous expression of MIK2 and MIK2 variants in *N. benthamiana*.** At least four independent biological replicates were performed (n≥4 plants), with each biological replicate represented by four technical replicates. Cumulative RLUs representing ROS production are shown after treatment with H_2_O (white) or the SCOOP peptide indicated (1 μM, grey). Significance was tested by performing a paired Wilcoxon rank-sum test.

Table S1: **Primers used in this study.**

| **Gene** | **Variant** | **Primer orientation** | **Sequence** |
| --- | --- | --- | --- |
| *AT4G08850* | D246G | fw | GTCTAGgTAGGAACAACCTTACCGGTAAAATCCCTT |
| *AT4G08850* | D246G | rev | TTGTTCCTAcCTAGACATAGCTCTCTAAGGTTGGGTA |
| *AT4G08850* | N268G | fw | TCTTCTCggTATGTTTGAAAATCAGCTCTCTGGTG |
| *AT4G08850* | N268G | rev | AACATAccGAGAAGAGTTACATTCTTCAAATTCCCGAA |
| *AT4G08850* | S292G | fw | TACACTTgGTCTCCACACAAATAAGCTTACCGGT |
| *AT4G08850* | S292G | rev | GGAGACcAAGTGTATCTAAAGCGGTCATATTACCAA |
| *AT4G08850* | H294G | fw | GTCTCggCACAAATAAGCTTACCGGTCCAATAC |
| *AT4G08850* | H294G | rev | TTTGTGccGAGACTAAGTGTATCTAAAGCGGTCATA |
| *AT4G08850* | H316G | fw | CGTTCTTggTCTTTACCTGAATCAACTCAATGGTTC |
| *AT4G08850* | H316G | rev | TAAAGAccAAGAACGGCTAGGGTTTTGATGTTTCC |

Dataset S1 (separate file): **Overview SCOOP mining.**

Dataset S2 (separate file): **Overview of mined assemblies for locus analyses, overview of Arabidopsis anchor genes *PROSCOOP* loci, and overview of contiguous *INR* loci (+ coordinates) and MIK2-orthologues.**

Dataset S3: **Plasmid maps of constructs used in this study.**

Dataset S4: **AFM predicted structures (.pdb) and confidence metrics (.pae)**

Dataset S5: **Unedited files (.tiff) of co-IP and western blotting.**
