## Supplementary figures and images for "Leveraging co-evolutionary insights and AI-based structural modeling to unravel receptor-peptide ligand-binding mechanisms"

### Co-IP_anti-bak1_3.tif

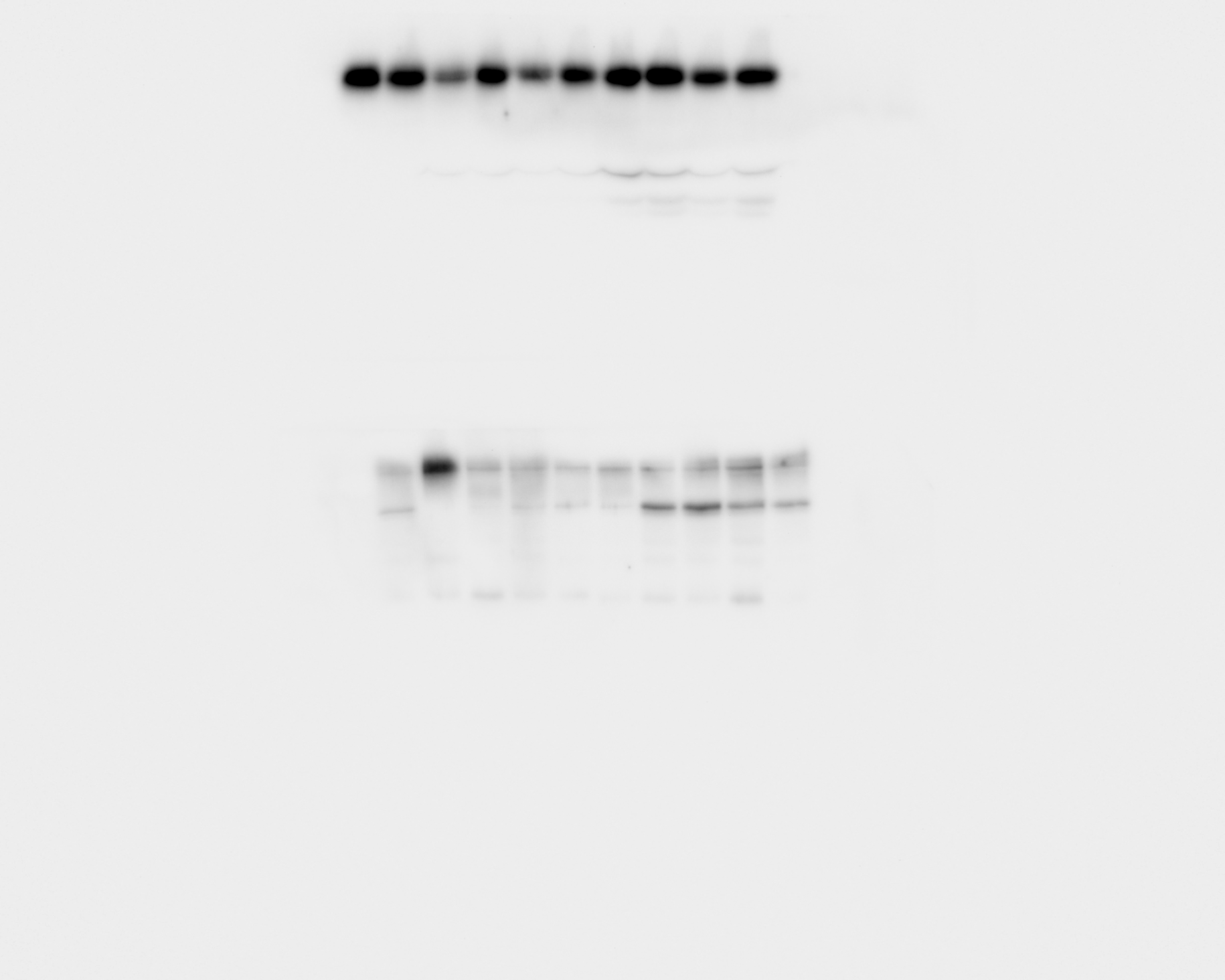

### Co-IP_anti-gfp mik2 inputs_7.tif

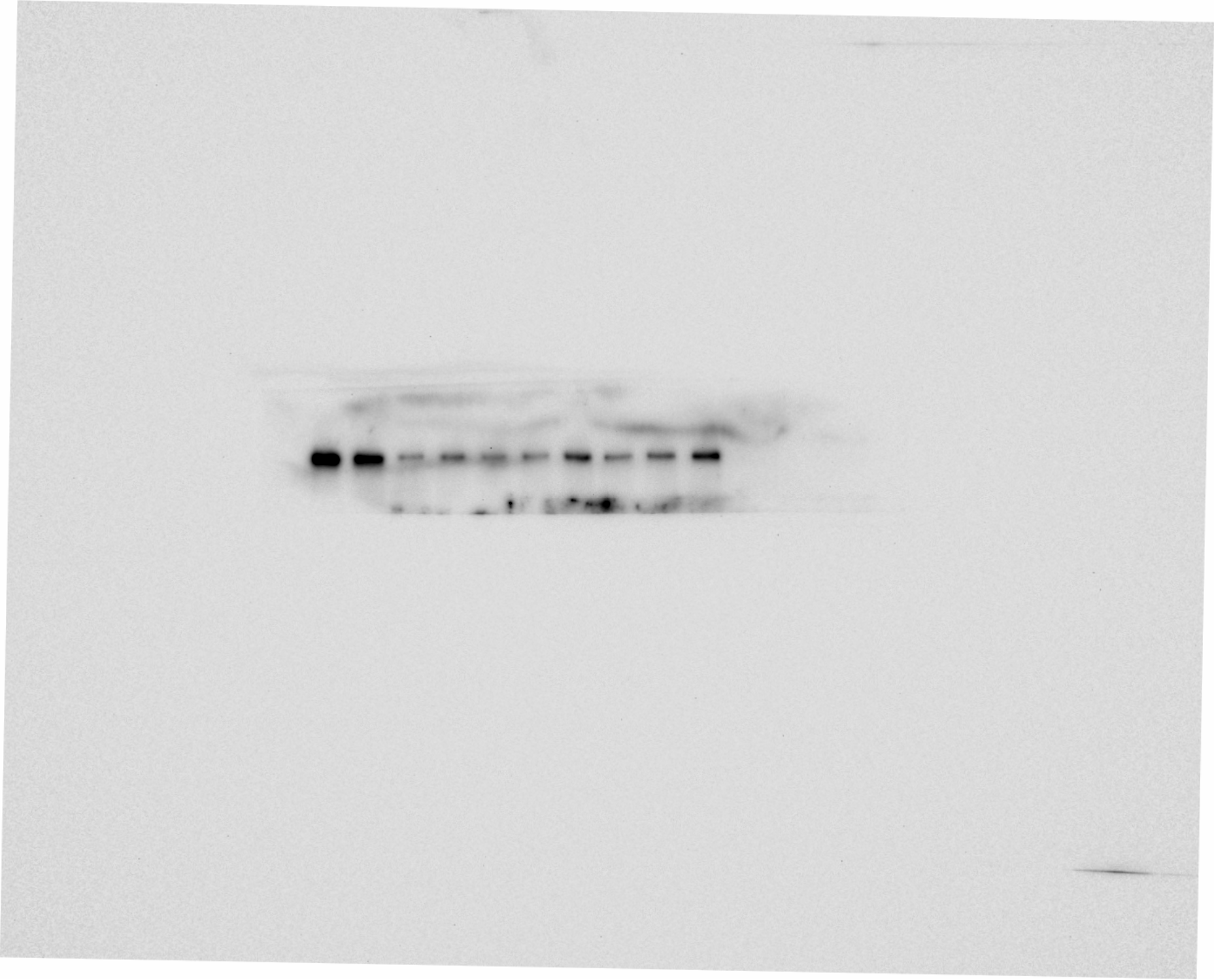

### Co-IP_anti-gfp mik2 ips_6_adjusted.tif

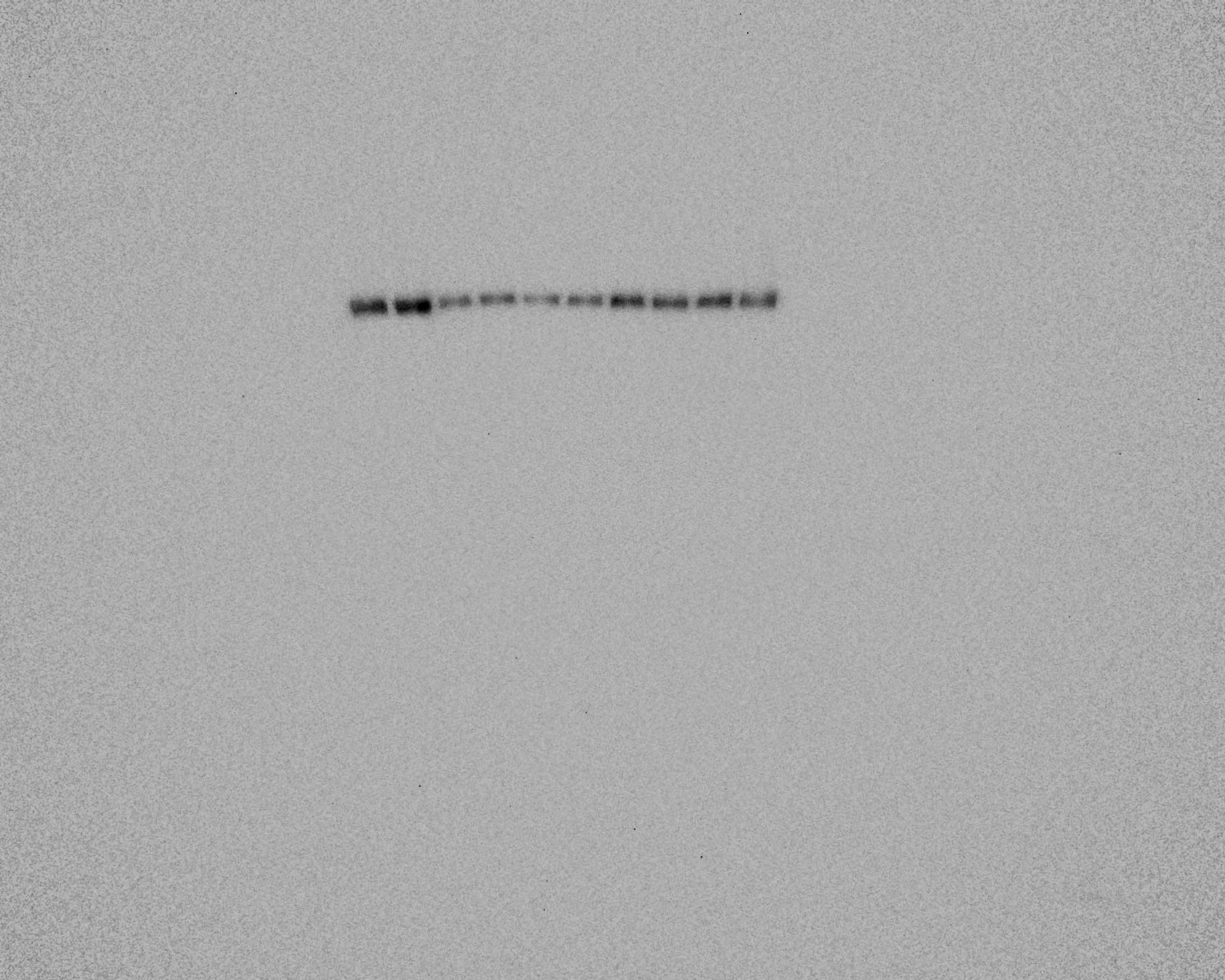

### Co-IP_cbbg250.tif

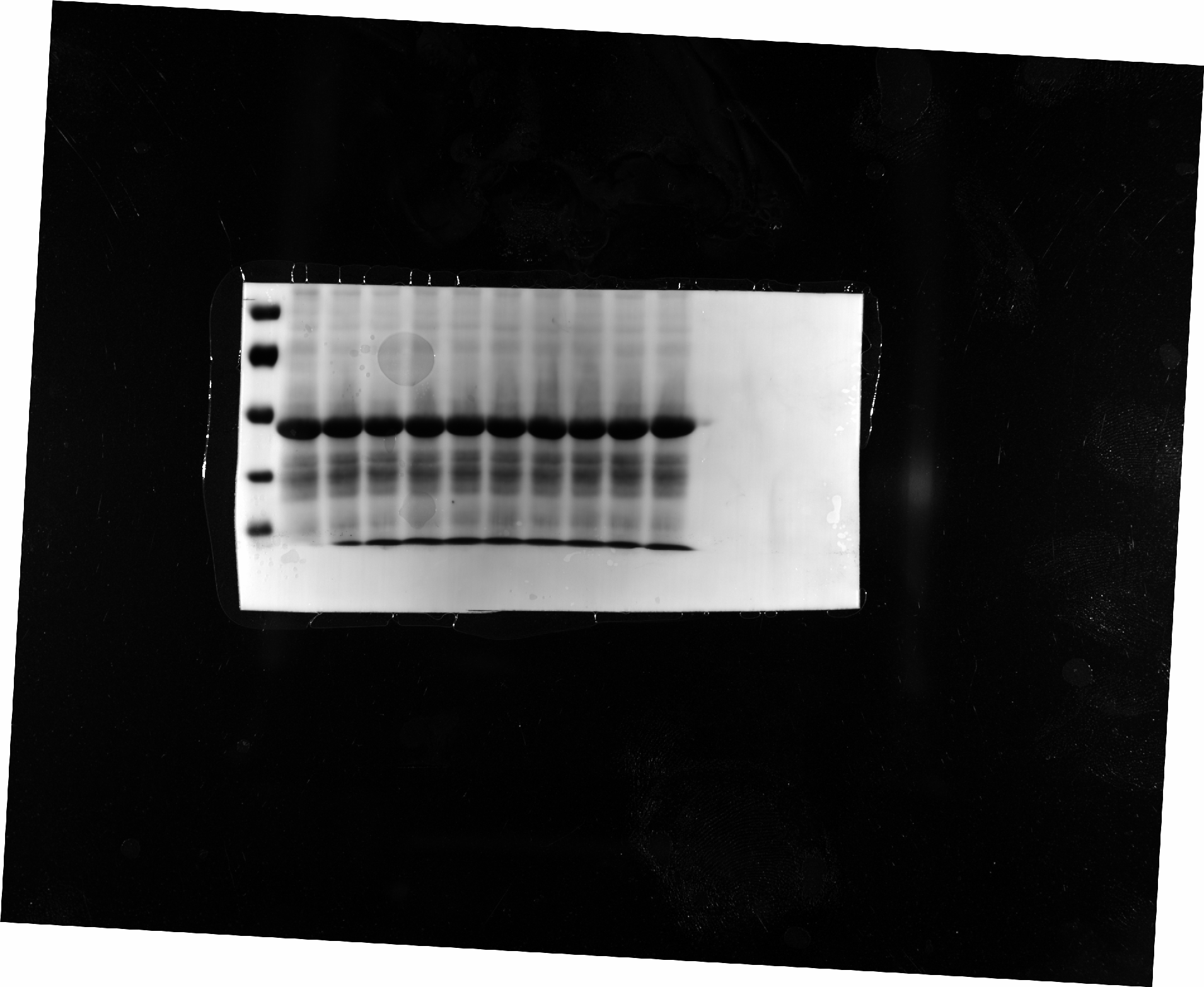

### extracellularMIK2SCOOP2_1806d_pae.png

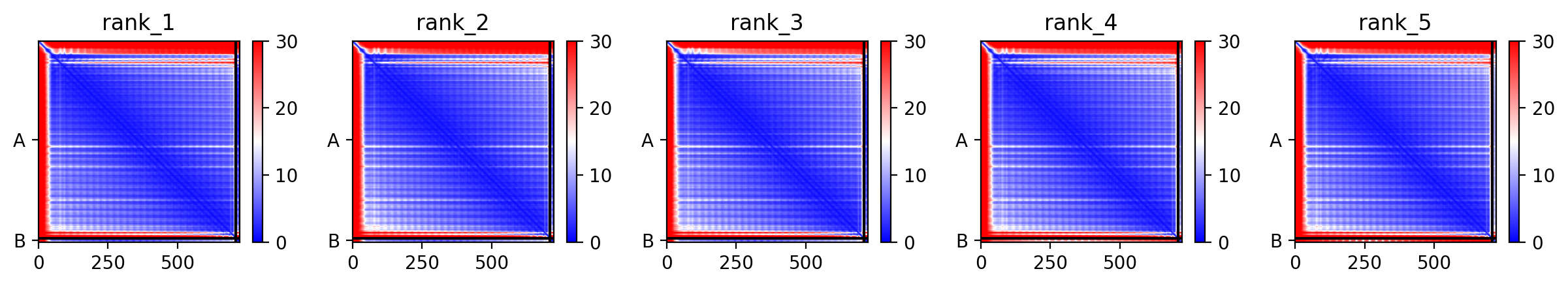

### extracellularMIK2SCOOP5_43154_pae.png

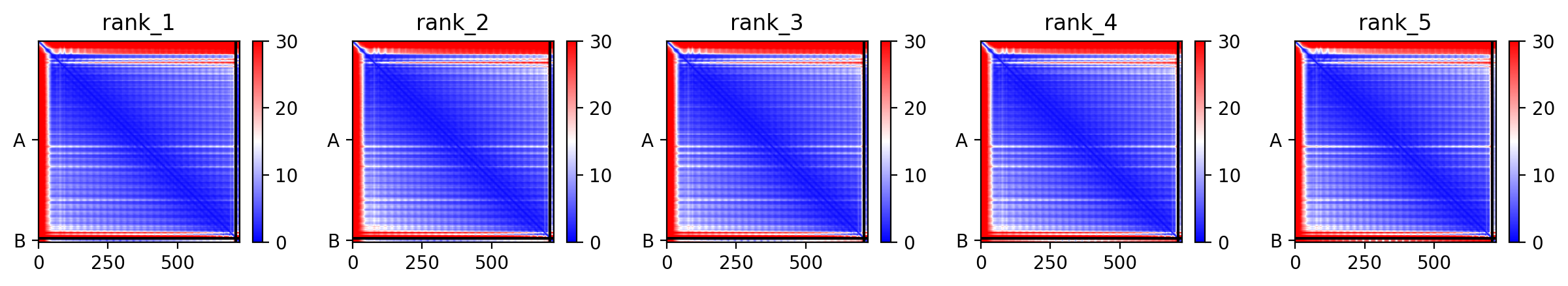

### extracellularMIK2SCOOP7_6c0d1_pae.png

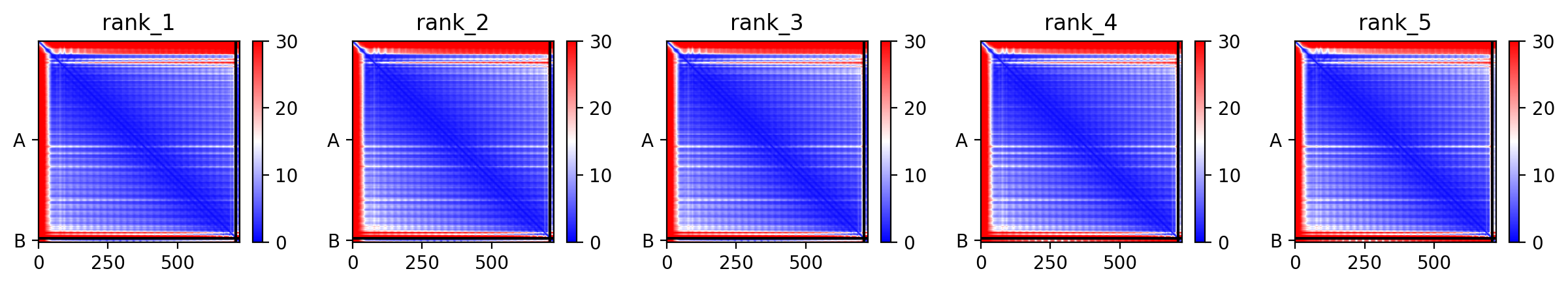

### extracellularMIK2SCOOP12_0e359_pae.png

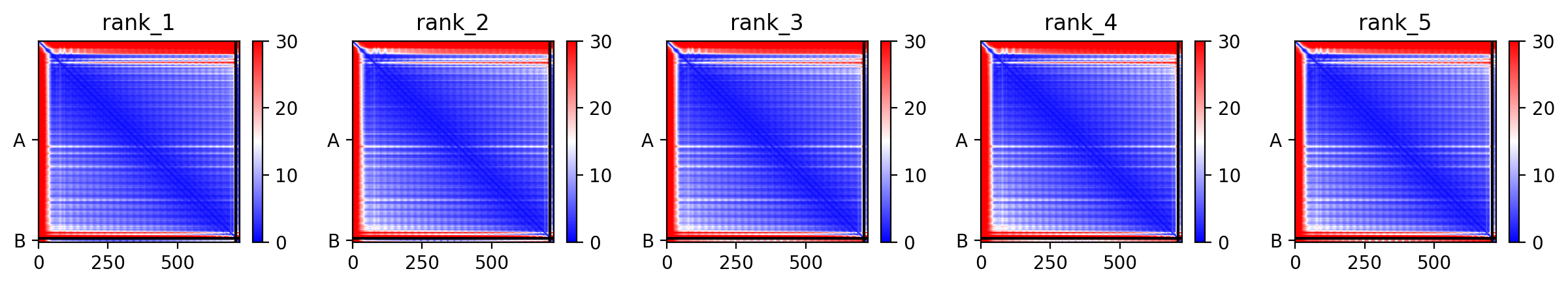

### extracellularMIK2SCOOP17_0b7e7_pae.png

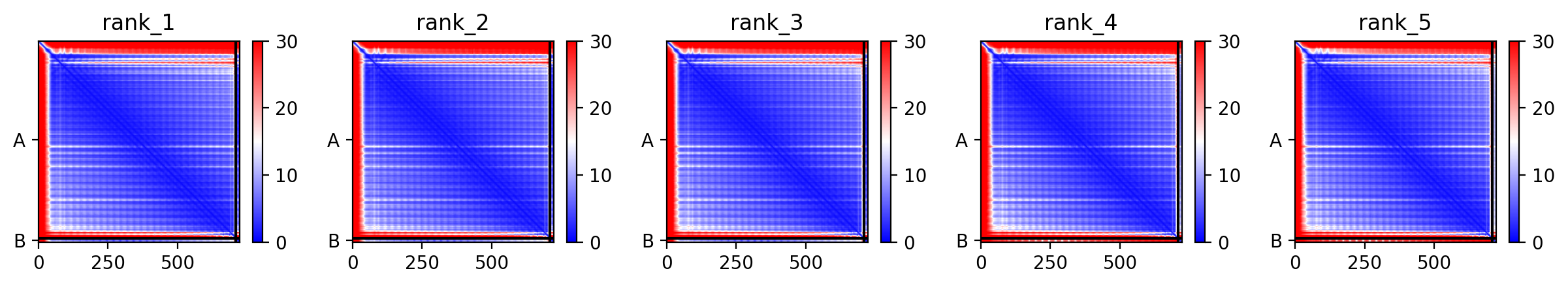

### extracellularMIK2SCOOP21_48952_pae.png

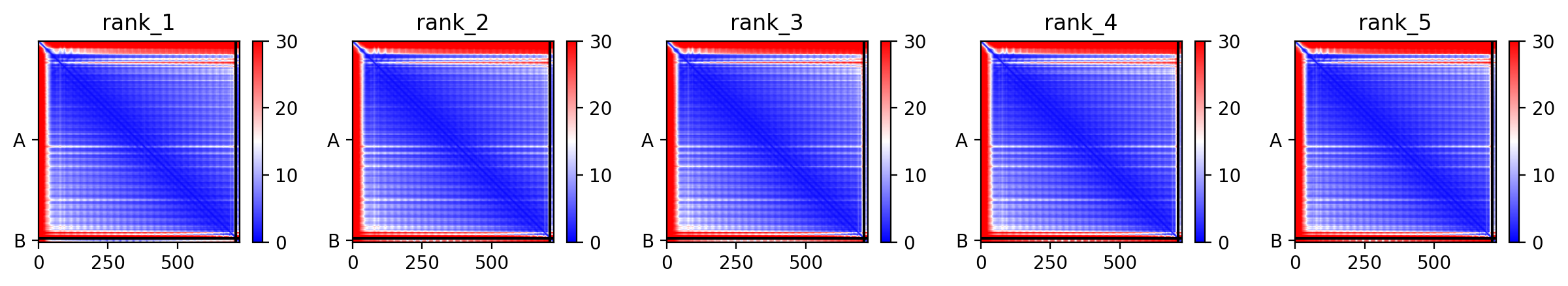

### extracellularMIK2SCOOP22_d2e56_pae.png

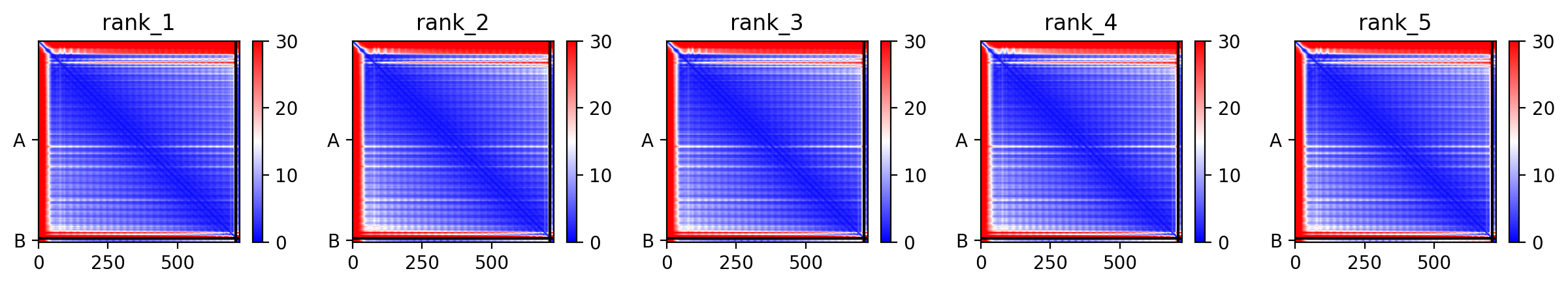

### extracellularMIK2SCOOP23_99635_pae.png

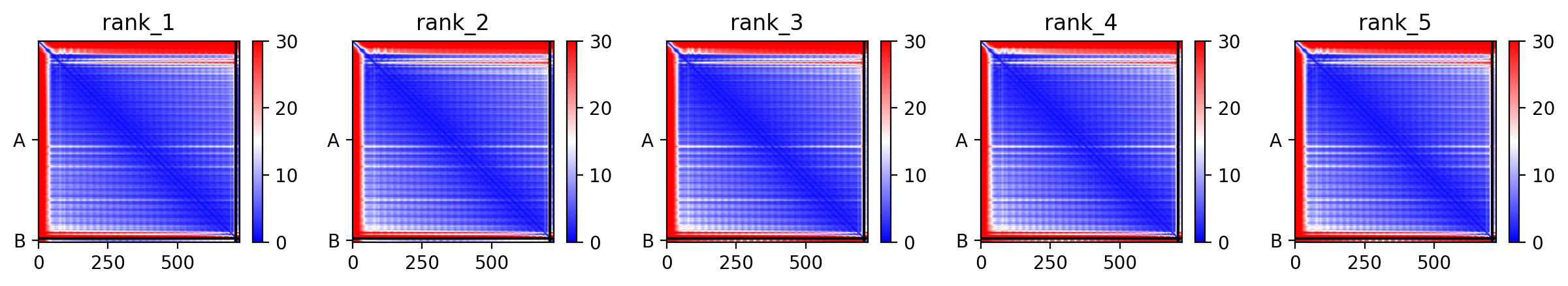

### extracellularMIK2SCOOP24_f83a9_pae.png

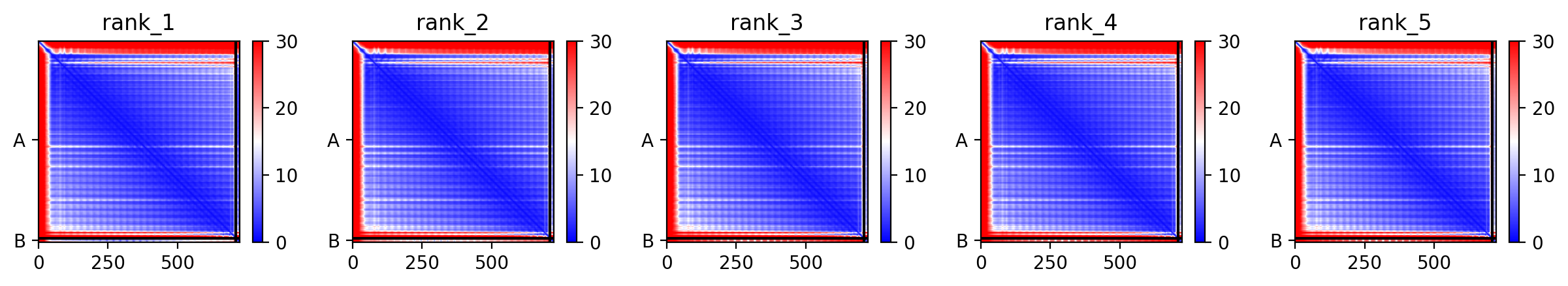

### extracellularMIK2SCOOP41_13_f0a92_pae.png

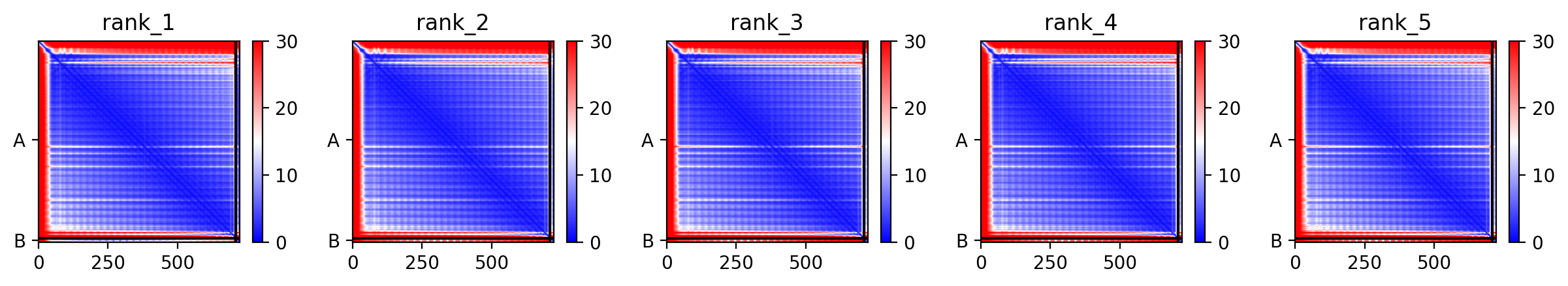

### extracellularMIK2SCOOP46_pae.png

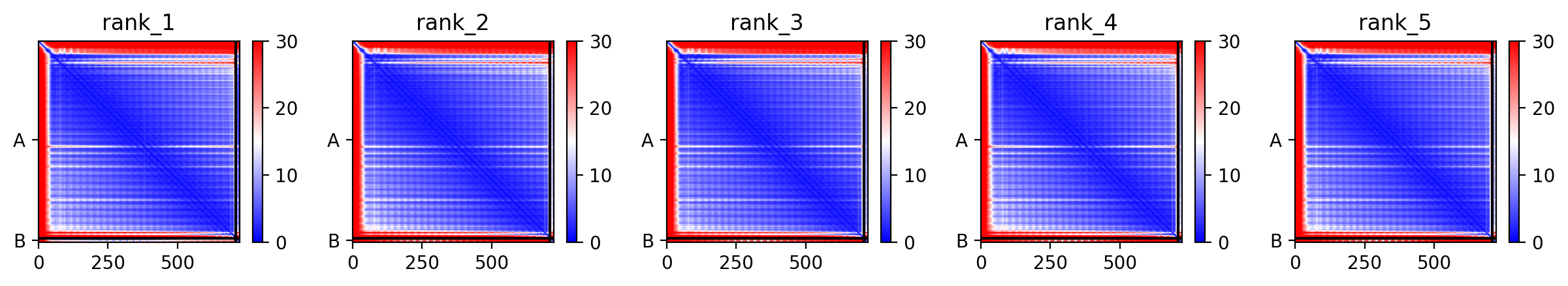

### extracellularMIK2SCOOP47_MIK2-47_pae.png

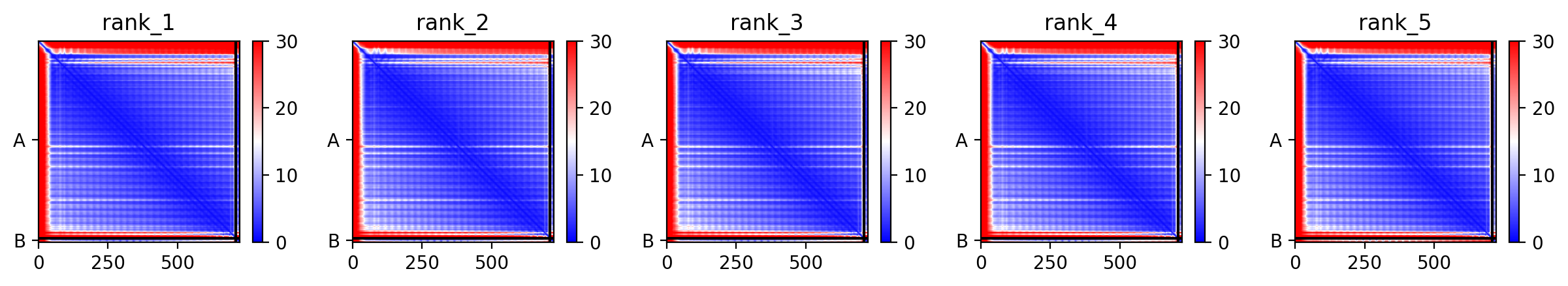

### Western_user 2023-09-19 08h55m30s(Composite).tif

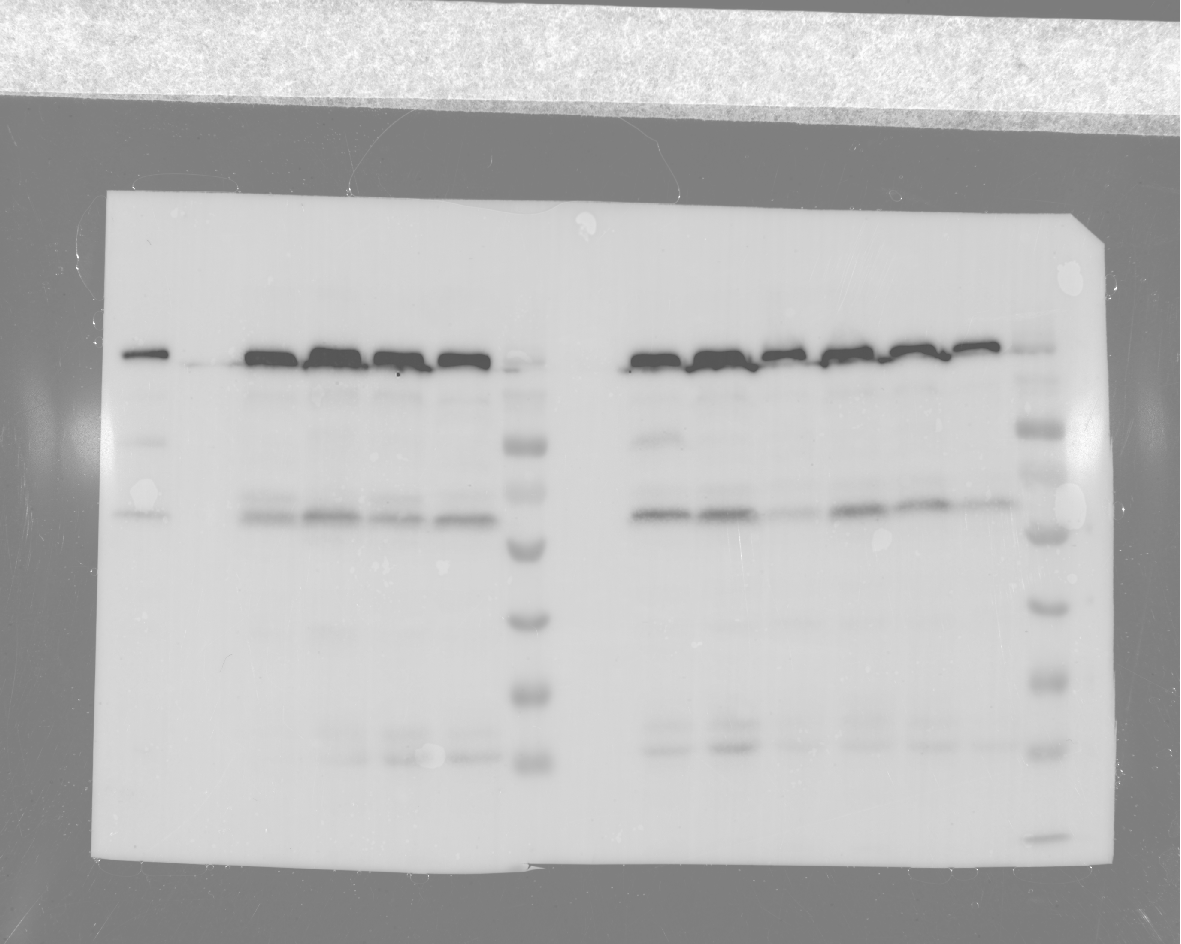

### Western_user 2023-09-19 09h05m43s(Colorimetric).tif

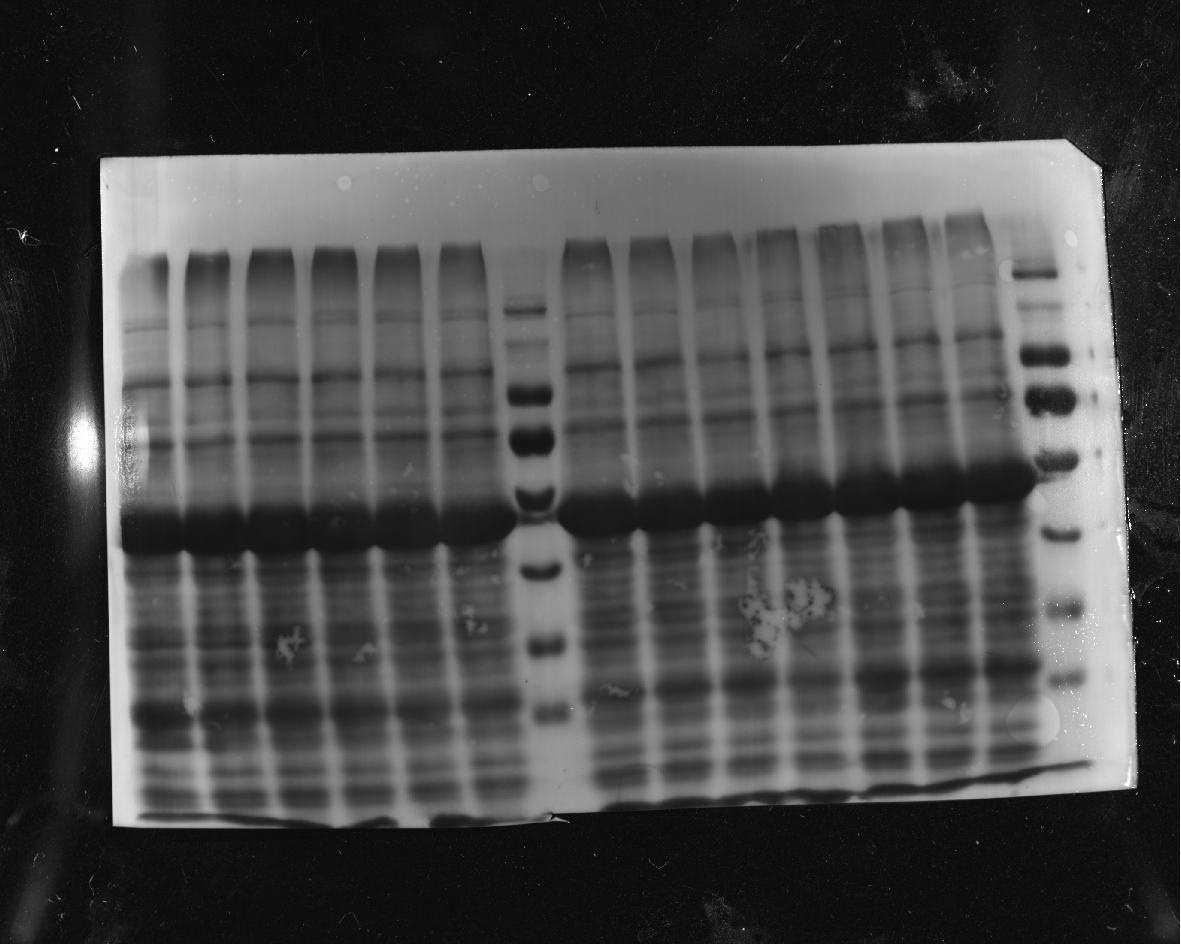
